# Longitudinal sex-at-birth and age analyses of cortical structure in the ABCD Study^®^

**DOI:** 10.1101/2024.06.10.598367

**Authors:** Andrew T. Marshall, Shana Adise, Eric C. Kan, Elizabeth R. Sowell

## Abstract

While the brain continues to develop during adolescence, such development may depend on sex-at-birth. However, the elucidation of such differences may be hindered by analytical decisions (e.g., covariate selection to address body/brain-size differences) and the typical reporting of cross-sectional data. To further evaluate adolescent cortical development, we analyzed data from the Adolescent Brain Cognitive Development Study^SM^, whose cohort of 11,000+ youth participants with biannual neuroimaging data collection can facilitate understanding neuroanatomical change during a critical developmental window. Doubly considering individual differences in the context of group-level effects, we analyzed regional changes in cortical thickness, sulcal depth, surface area, and volume between two timepoints (∼2 years apart) in 9-to 12-year-olds assigned male or female sex-at-birth. First, we conducted linear mixed-effects models to gauge how controlling for intracranial volume, whole-brain volume (WBV), or a summary metric (e.g., mean cortical thickness) influenced interpretations of age-dependent cortical change. Next, we evaluated the relative changes in thickness and surface area as a function of sex-at-birth and age. Here, we showed that WBV (thickness, sulcal depth, volume) and total cortical surface area were more optimal covariates; controlling for different covariates would have substantially altered our interpretations of overall and sex-at-birth-specific neuroanatomical development. Further, we provided evidence to suggest that aggregate change in how cortical thickness is changing relative to surface area is generally comparable across those assigned male or female sex-at-birth, with corresponding change happening at slightly older ages in those assigned male sex-at-birth. Overall, these results help elucidate neuroanatomical developmental trajectories in early adolescence.

**Significance Statement:** While most of our brain’s development happens early in life, much of it still happens in adolescence. Because many factors can alter those developmental trajectories, it is important to evaluate the shape/timing of those trajectories (i.e., what generally constitutes typical brain development). Here, we showed that our understanding of those trajectories can be affected by how we choose to analyze them. First, we showed that the way researchers address differences in brain/body size affects how we interpret regional variation in brain change over time. Further, we showed that it is important to consider how similar patterns of development may simply be happening at different ages in different groups. These results support a relatively novel way of analyzing adolescent brain development.

## Introduction

Research has consistently reported age-related neuroanatomical changes during critical windows of development, like adolescence (e.g., Giedd et al., 1999; Sowell et al., 2004; Tamnes et al., 2017; Tamnes et al., 2010; Wierenga et al., 2014). Therein, the brain is subject to substantial cortical maturation, such as regional synaptic pruning (Petanjek et al., 2011), gray-matter thinning (Gogtay et al., 2004; Raznahan et al., 2014; Sowell et al., 1999), and increased white-matter myelination of neuronal connections (Asato et al., 2010; Benes, 1989; Simmonds et al., 2014). Generally, cortical expansion (increased surface area) parallels its concurrent thinning, even though other research has reported regionally specific cortical thickening and/or contraction during adolescence (e.g., Mills et al., 2021; Schnack et al., 2015; Sowell et al., 2004; Tamnes et al., 2017; Vijayakumar et al., 2016).

Notably, cortical development may also be sex-specific (e.g., Cosgrove et al., 2007; Duerden et al., 2020; Gennatas et al., 2017; Herting et al., 2015; Lenroot & Giedd, 2010; Ruigrok et al., 2014; Sowell et al., 2007; Torgerson et al., 2024). For example, participants who are female had thicker cortices than those who are male, with there being regionally specific, nonlinear sex-by-age interactions from 7-to 87-years-old (Sowell et al., 2004). In 11- to 20-year-olds, participants who are male showed larger, multi-regional cortical surface areas and volumes relative to participants who are female, but there were fewer sex-specific differences in how cortical thickness, surface area, and volume changed with age (Vijayakumar et al., 2016). Similarly, in 8- to 22-year-olds, there were reductions in cortical volume with age, but these changes did not differ by sex (Tamnes et al., 2013). Interesting, in 4.5- to 18-year-olds, developmental trajectories of cortical and subcortical volume were more curvilinear in participants who are female and more linear in those who are male (Brain Development Cooperative Group, 2012). Despite the plethora of research producing differing results, one robust finding has been larger brain size in participants who are male versus female (see Bhargava et al., 2021). However, the overall inconsistency of such results warrants continued analyses of these phenomena to understand the trajectories of (and deviations from) neurotypical development.

The seminal Adolescent Brain Cognitive Development Study^SM^ (ABCD Study®, hereafter “ABCD”) offers a unique opportunity to elucidate such discrepancies. Indeed, one of its main goals is to “develop national standards for normal brain development in youth, by defining the range and pattern of variability in trajectories of brain development” (Jernigan, Brown, & ABCD Consortium Coordinators, 2018, p. 2). Here, in ABCD participants, via two analytical approaches which simultaneously considered cortical development at the group/population level and individual level. we characterized differences in age-related neurodevelopment between two timepoints (∼2 years apart) in 9- to 12-year-olds assigned male or female sex-at-birth.

First, as body/brain size may account for sex-specific differences in cortical structure (Mills et al., 2016; Pintzka et al., 2015), we determined how population-level regression coefficients relating age to regional cortical structure depended on which whole-brain metric was included in our models (e.g., whole-brain volume, intracranial volume), with our goal being to covary for the metric that more precisely captured population- and individual-level developmental change. Secondly, as there have been relatively few studies on how different cortical measures covary with age (Norbom et al., 2021), we evaluated how individual-level annual percentage changes (APC) of cortical thickness versus surface area varied by region and sex-at-birth. Given the bivariate nature of these analyses, we employed a relatively novel approach to describe the trajectory of cortical development (see Schnack et al., 2015), one which captured the changing relationships between the APCs of cortical thickness and surface area as rotating vectors around a coordinate plane. Overall, by considering individual differences (and their distributions) within the larger ABCD sample, we have provided another piece to the puzzle to elucidate the possibility of sex-at-birth differences in neuroanatomical development during adolescence.

## Methods

### Participants

ABCD is a 10-year longitudinal study at 21 U.S. study sites (Jernigan, Brown, & Dowling, 2018). Primarily using school-based enrollment (Garavan et al., 2018), the consortium enrolled nearly 11,900 9- and 10-year-old youth. Our data came from the 2021 ABCD 4.0 public data release (doi: 10.15154/1523041), which included baseline data for 11,876 youth and two-year follow-up data for 10,414 youth (i.e., the two appointment years with neuroimaging data in the 4.0 release) (see Table 1). Centralized IRB approval was obtained from the University of California, San Diego. Study sites obtained approval from their local IRBs. Parents provided written informed consent and permission; youth provided written assent. Data collection and analysis complied with all ethical regulations.

**Table 1.** Participant Demographics.

|  | ABCD Cohort –<br>Baseline (%) | Sample in This<br>Report (%) |
| --- | --- | --- |
| **Youth Sex** |  |  |
| Male | 6,196 (52.2%) | 3,412 (54.0%) |
| Female | 5,680 (47.8%) | 2,911 (46.0%) |
| **Annual Household Income** |  |  |
| <\$50K (Low) | 3,223 (27.1%) | 1,642 (26.0%) |
| \$50-100K (Mid) | 3,071 (25.9%) | 1,748 (27.6%) |
| >\$100K (High) | 4,564 (38.4%) | 2,438 (38.6%) |
| Missing/Undefined | 1,018 (8.6%) | 495 (7.8%) |
| **Caregiver Education** |  |  |
| ≤12th Grade/No Diploma | 593 (5.0%) | 286 (4.5%) |
| High-school Graduate/GED or<br>Equivalent | 1,132 (9.5%) | 536 (8.5%) |
| Some College, No<br>Degree/Associate's Degree | 3,079 (25.9%) | 1,617 (25.6%) |
| Bachelor's Degree | 3,015 (25.4%) | 1,652 (26.1%) |
| Master's/Professional Degree,<br>Doctorate | 4,043 (34.0%) | 2,223 (35.2%) |
| Missing/Undefined | § (§%) | § (§%) |
| **Youth Race** |  |  |
| American Indian / Alaska Native | 62 (0.5%) | 35 (0.6%) |
| Asian | 275 (2.3%) | 143 (2.3%) |
| Black | 1,869 (15.7%) | 856 (13.5%) |
| Native Hawaiian / Pacific<br>Islander | § (§%) | § (§%) |
| Other | 1,959 (16.5%) | 1,040 (16.4%) |
| White | 7,524 (63.4%) | 4,160 (65.8%) |
| Missing/Undefined | 171 (1.4%) | 80 (1.3%) |
| **Youth Ethnicity** |  |  |
| Hispanic | 2,411 (20.3%) | 1,282 (20.3%) |
| Not Hispanic | 9,312 (78.4%) | 4,966 (78.5%) |
| Missing/Undefined | 153 (1.3%) | 75 (1.2%) |
| **Total** | **11,876 (100%)** | **6,323 (100%)** |

*Note.* The data in the “ABCD Cohort” column refer to those derived from the 4.0 ABCD Study Data Release (October 2021). The “Other” Race/Ethnicity category includes those who identified “Other Race” and those who identified more than one race (i.e., multiracial). Values of small cell sizes ( $n < 20$ ) were suppressed (i.e., noted by §).

### Neuroimaging

ABCD neuroimaging data collection and processing procedures have been described in detail (Hagler et al., 2019). From the publicly available data, we analyzed four measures of regional cortical brain structure (derived using FreeSurfer v.7.1.1 Desikan-Killiany atlas on acquired T_1_w MRI volumes): thickness, surface area, volume, and sulcal depth (Dale et al., 1999; Fischl & Dale, 2000). The Desikan-Killiany atlas includes 34 bilateral regions (68 regions total). Whole-brain volume and intracranial volume were included in analyses. Depending on the ABCD study site, Siemens, Philips, and GE scanners were used. All neuroimaging parameters are available: https://abcdstudy.org/images/Protocol_Imaging_Sequences.pdf. T_1_w images were corrected for gradient nonlinearity distortions per scanner-manufacturer guidelines, with further details comprehensively described previously (Hagler et al., 2019). ABCD data are publicly available on the NIMH Data Archive (https://data-archive.nimh.nih.gov/abcd).

### Statistical Analyses

Per recommendations within ABCD’s 4.0 data-release notes, the neuroimaging data from 8 youth participants were excluded (i.e., due to event mislabeling or sMRI processing failures), as were 209 participants whose T1w data were not recommended for analysis by ABCD’s Data Analysis, Informatics, & Resource Center (remaining *n* = 11,659). With our goal being to look at within-individual change, we also removed 4,227 participants who did not have both baseline and two-year follow-up structural MRI data (remaining *n* = 7,432). Lastly, as the ABCD cohort includes siblings, we chose to control for family relatedness by only including singletons or one sibling per family in analysis; here, using MATLAB’s *datasample* function (seed = 1), we randomly selected one sibling from the corresponding family groups (remaining *n* = 6,323) [MATLAB Version: 9.13.0.2126072 (R2022b) Update 3]. The demographics of the entire ABCD cohort at baseline versus those of the current analyzed sample (*n* = 6,323) are shown in Table 1.

### Sex × Age Interactions on Cortical Structure

#### Whole-brain covariate selection

First, we evaluated how controlling for different types of whole-brain metrics affected the fixed-effects regression coefficients for associations between age and regional cortical structure. For each of the 4 cortical metrics across 68 bilateral regions, we ran 4 linear mixed-effects models:

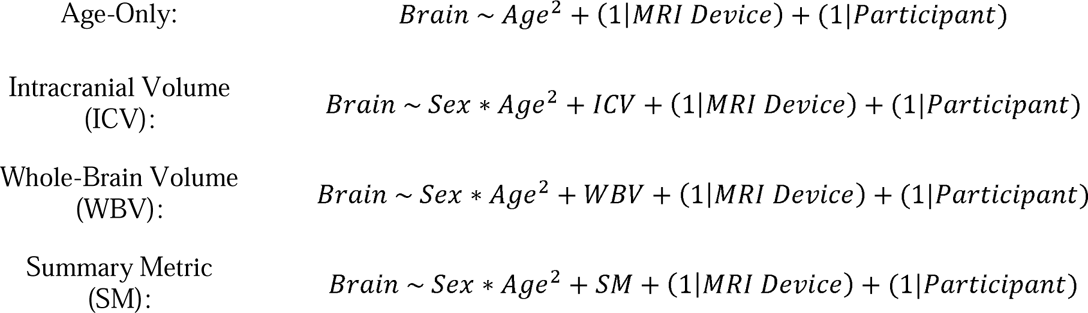

Here, age (in months), ICV, WBV, and SM were continuous, mean-centered predictors. Sex was an effects-coded categorical factor with male/female (M/F) coded as -1/+1 (i.e., when referring to sex-at-birth in the context of the regression-model terms, references to the factor “Sex” refers to sex-at-birth). By-MRI device and by-participant intercepts were included as random effects. All models used a random initial value for iterative optimization. SM was dependent on the corresponding metric (i.e., mean cortical thickness, mean sulcal depth, total area, or total volume).

For each analysis (i.e., region-by-region), participants’ data were considered outliers (and excluded from analysis) if either of their datapoints (baseline or two-year follow-up) exceeded more than three scaled median absolute deviations away from the median (Leys et al., 2013). This resulted in between 15 and 301 participants being removed from cortical thickness analyses (*n*’s = 6,022 to 6,308); between 11 and 212 participants from sulcal depth analyses (*n*’s = 6,111 to 6,312); between 6 and 127 participants from surface area analyses (*n*’s = 6,196 to 6,317); and, between 10 and 184 participants from volume analyses (*n*’s = 6,139 to 6,313). Because participants only had 2 datapoints in analysis, spanning approximately 2 years apart, we focused on how age was associated with brain structure (and how these associations were moderated by sex at birth); however, we included a quadratic fixed-effect term of *Age*^2^ in all models to capitalize on the entire cross-sectional age range of ABCD, given past research describing curvilinear changes in brain structure across longer stretches of adolescence (e.g., Mills et al., 2021).

To determine the more optimal whole-brain covariate for each of the thickness, sulcal depth, surface area, and volume analyses, we performed two analyses based on the supposition that (1) the age-only model should most accurately convey actual intraindividual changes with age and (2) a proper whole-brain control metric should not create estimates that deviate far from the age-only coefficient. First, we ran a simple correlation (Pearson’s *r*) between the fixed-effects coefficients from the age-only models of the 68 ROIs for each of thickness, sulcal depth, surface area, and volume data against those from the ICV, WBV, and SM models; a higher correlation reflected greater similarities between the two sets of coefficients. Second, we evaluated how well the age-only coefficients “fit” those from the ICV, WBV, and SM models, in that we computed the residual sum of squares between each of the ICV, WBV, and SM models and the age-only model (i.e., we summed the squared differences between the coefficients from each of the former and the latter); here, smaller values reflected greater similarities between two sets of coefficients.

#### Sex × Age interactions

After establishing a proper whole-brain covariate for each of cortical thickness, sulcal depth, surface area, and volume, we used the corresponding models to evaluate how sex-at-birth moderated associations between age and brain structure using either the ICV, WBV, or SM models. The corresponding effect sizes for age are represented by partial correlation coefficients (*r_p_*), which control for all model covariates and are calculated using the corresponding *t*-statistic and degrees of freedom (Nakagawa & Cuthill, 2007). The strengths of the Sex × Age interactions were calculated similarly (i.e., using the corresponding *t*-statistic and degrees of freedom). The 95% CIs of the effect sizes were derived from the sample variance of the partial correlation (Aloe & Thompson, 2013). The Benjamini-Hochberg (1995) false-discovery-rate (FDR) algorithm was used to correct for multiple comparisons.

### Annual Percentage Change

Our second set of analyses evaluated individual differences in age-associated structural change using an annual percentage change (APC) metric (https://www.cdc.gov/nchs/hus/sources-definitions/average-annual-rate-of-change.htm):

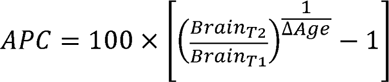

in which *Brain_T_*_2_ and *Brain_T_*_1_ refer to sMRI cortical metrics at the baseline (T1) and the two-year follow-up appointments (T2), respectively, and Δ*Age* refers to the change in age (in years; i.e., change in age in months divided by 12) between T1 and T2. We specifically analyzed the APC of cortical thickness and surface area to evaluate the differential rates of contraction versus expansion (surface area [CSA]) and thinning versus thickening (thickness [CT]). Participants’ APC data were considered outliers (and excluded from analysis) if, across the sample, they exceeded more than three scaled median absolute deviations away from the median (Leys et al., 2013).

#### Comparing APC_CT_ to APC_CSA_

For each region by sex-at-birth, one-sample *t*-tests were performed on APC_CT_ and APC_CSA_ to determine (1) if APC_CT_ and APC_CSA_ were significantly different from 0 and (2) the magnitude and direction of the APCs [thinning (-APC_CT_), thickening (+APC_CT_), contraction (-APC_CSA_), expansion (+APC_CSA_)]. Because there were both positive and negative APC values, we converted the aforementioned *t*-statistic to Cohen’s *d* (i.e., *t*-statistic divided by the square root of the degrees of freedom for the *t*-test); the absolute Cohen’s *d* values (i.e., |*d*|) were then used to compare absolute rates of thickening/thinning and expansion/contraction. The presence of statistically significant differences in rates of thickening/thinning versus expansion/contraction for each region was identified based on whether there was any overlap in the corresponding 84% confidence intervals (CI) of the Cohen’s *d* values (see Austin & Hux, 2002; MacGregor-Fors & Payton, 2013). Here, the 84% CIs of the Cohen’s *d* values were computed as *d* ± *SEM* × *z*_0.92_, with:

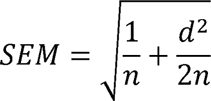

Therefore, this analysis allowed us to categorize whether each cortical region was, at the group level, thickening/thinning (cortical thickness) at faster (or slower) rates than it was expanding/contracting (cortical surface area), resulting in 17 possible categories (Table 2). Post hoc sensitivity analyses were also conducted with only individuals scanned by the same MRI machine at both timepoints (i.e., to verify whether the corresponding patterns were not solely due to any differences between MRI scanners).

**Table 2.**
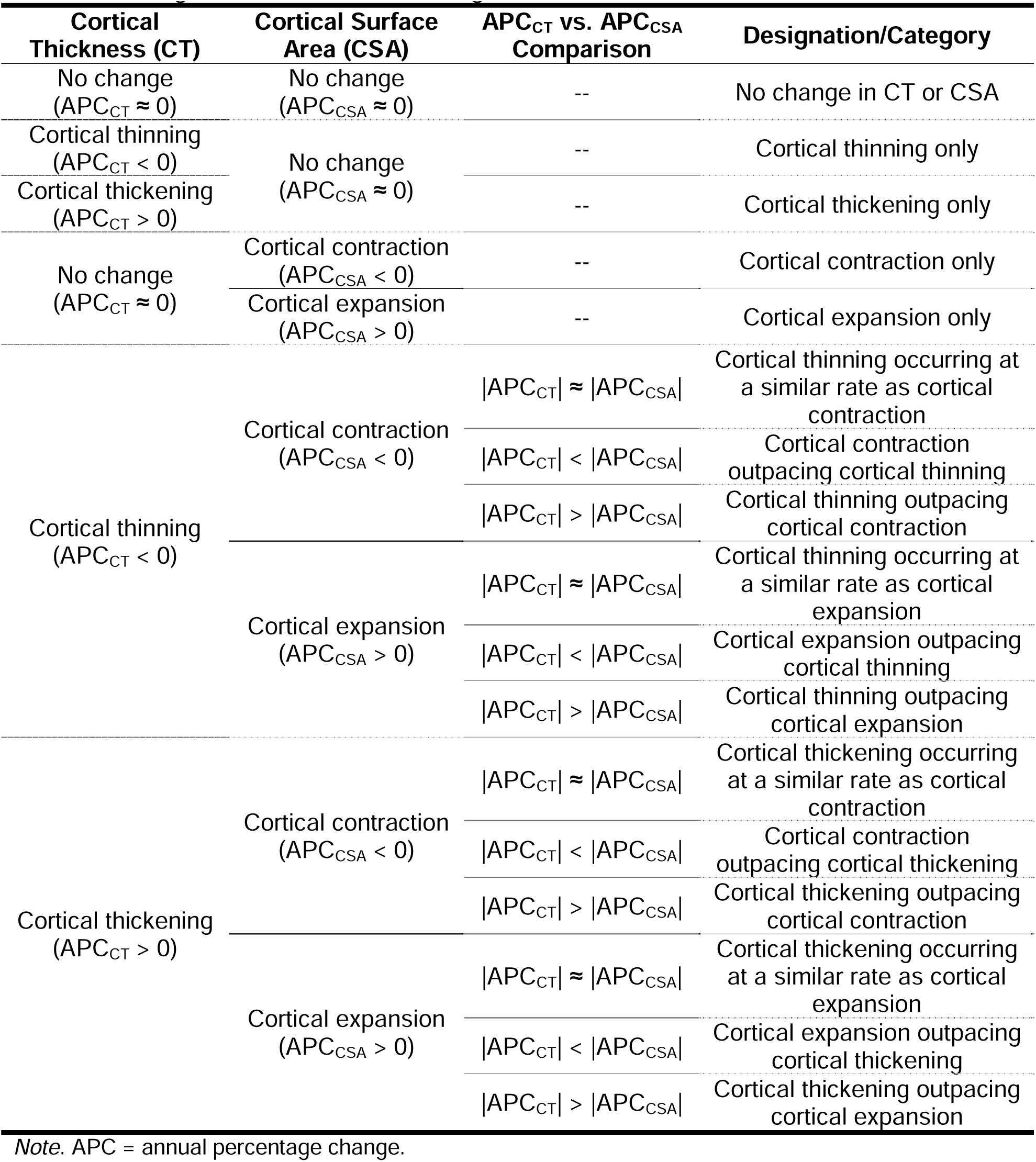
Categorization of cortical change.

#### Group-level representativeness

To evaluate how well each region’s APC_CT_-APC_CSA_ category (at the group level) represented the individuals in that group, we computed a representativeness summary proxy based on direction and strength of each individual’s APC_CT_ and APC_CSA_. For example, if one region (at the group level) showed cortical thinning as happening at a faster rate than cortical expansion, we determined whether each individual’s (1) APC_CT_ was less than zero, (2) APC_CSA_ was greater than zero, and (3) whether the absolute value of APC_CT_ was greater than the absolute value of APC_CSA_; thus, by way of this example, there was a 12.5% chance that each individuals’ data would randomly meet these three criteria. For regions in which only the group-level APC_CSA_ or APC_CT_ was significantly different from 0, we checked whether the individuals’ change-over-time data matched the direction of that group-level APC (i.e., a 50% chance that individuals’ data would randomly meet that one criterion). For regions in which there were similar rates of thickening/thinning (APC_CT_) versus expanding/contracting (APC_CSA_), we checked whether the absolute values of APC_CT_ and APC_CSA_ did not differ by more than 2%, which was the greater of the approximate standard deviations of APC_CT_ and APC_CSA_ (APC_CT_ = 1.45, APC_CSA_ = 1.98) (see Zhou et al., 2015). After calculating the proportion of individuals who showed the developmental trajectories characterizing the group, we determined whether the CI of that proportion (by sex-at-birth and brain region) also included the chance proportion, in order to gauge how well the groups’ individuals were represented by the group-level APC_CT_-APC_CSA_ category. Again, post hoc sensitivity analyses were run only with individuals scanned by the same MRI machine at both timepoints.

#### Circular distribution and analysis

Next, we evaluated the distribution of sex-at-birth differences in the relative APC_CT_-APC_CSA_ trajectories for individuals scanned by the same machine at both timepoints. Given an individual-differences scatterplot that can be generated from these data (x-coordinate = APC_CT_, y-coordinate = APC_CSA_), we converted each individual’s x-y coordinates to radians via a two-argument arctangent function *atan2*. Intuitively, this derived radian value (for each individual participant) represents a direction (or vector) on the coordinate plane referring to how much the individual’s regional cortical thickness is changing relative to how much that individual’s regional cortical surface area is changing. Accordingly, we treated each direction as data points on a circular distribution around the origin of the coordinate plane.

To evaluate the amount of variance explained by group (i.e., sex-at-birth) in these APC_CT_-APC_CSA_ vectors, we conducted one-way analyses of variance (ANOVA) on these circular data for each bilateral region using the circular-statistics package *circular* (Agostinelli & Lund, 2023) in R (R Core Team, 2020). The Benjamini-Hochberg algorithm was used to correct for multiple comparisons. Regardless of the statistical significance of the ANOVA, we determined the variance explained by sex-at-birth using the sums of squares (SS) from the ANOVA output table (i.e., SS_Between_/SS_Total_). A total of 340,528 datapoints were included in these analyses, with each bilateral analysis included between 4,654 and 5,193 individuals. For ease of plotting, we partitioned individuals into 8 crude bins (i.e., thickening/thinning is happening faster/slower than expansion/contraction) and plotted polar histograms by cortical region and sex-at-birth.

#### APC_CT_-APC_CSA_ trajectories by age

We then evaluated how these circular, sex-at-birth-specific distributions may be influenced by age. Specifically, we calculated the mean angle (or direction) (in radians) by sex-at-birth, age, and ROI. Participant age was defined as the halfway point between the ages at the baseline and two-year follow-up appointments, and, for simplicity, was collapsed into 13 levels approximately characterized by two-month ranges. Given the radian values for APC_CT_-APC_CSA_ described above, the mean angles (or directions) by age, sex-at-birth, and ROI of the APC_CT_-APC_CSA_ circular distributions were computed as:

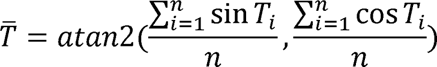

in which *T̄* is the mean direction (or angle), *T_i_* is individual *i*’s radian value, and *n* is the number of individuals. Intuitively, *T̄* is the average angle (or direction) on the coordinate plane referring to the rate of change of cortical thickness (APC_CT_) relative to that of cortical surface area (APC_CSA_) (e.g., at approximately 120 months of age, how much is the cortex thinning relative to how much is it expanding?).

Lastly, the 95% CIs for the mean directions (or angles) were computed as described previously (Mardia & Jupp, 2000, p. 124); because 2.48% of these computed 95% CIs were complex numbers, we used the absolute value of the CIs to calculate the lower and upper CI bounds. In order to plot the mean directions (or angles) *T̄* on a standard x-y plane with age on the abscissa and *T̄* on the ordinate, MATLAB’s *modulo* function was used to convert *T̄* to an inverted [0, 2π] scale. For example, *T̄* values equal to 3.14 (or π) mean that that brain region “Cortical thinning only” per Table 2). Further, for example, *T̄* values of 0 (or 2π) indicate that a showed cortical thinning (APC_CT_ < 0) with no change in cortical surface area (APC_CSA_ = 0) (i.e., region showed cortical thickening (APC_CT_ > 0) with no change in cortical surface area (APC_CSA_ = 0) (i.e., “Cortical thickening only” per Table 2). Accordingly, *T̄* values between 1.57 (π/2) and (APC_CSA_ < 0); *T̄* values between 3.14 (π) and 4.71 (3π/2) indicate that a region showed cortical 3.14 (π) indicate that a region showed cortical thinning (APC_CT_ < 0) and cortical contraction thinning (APC_CT_ < 0) and cortical expansion (APC_CSA_ > 0).

## Results

### Sample Characteristics

At baseline, youth participants were ∼9.9 years old (*SD* = 0.6; range = 8.9-11.0 years old). At the year-two follow-up appointment, youth participants were ∼11.9 years old (SD = 0.6; range = 10.6-13.8 years old). Compared to the entire ABCD cohort, our current sample was slightly more likely to have higher incomes and education levels (Table 1).

### Sex **×** Age Interactions on Brain Structure

#### Whole-brain covariate selection

First, we evaluated how controlling for different whole-brain metrics affected the fixed-effects coefficients of age at the individual level (Extended Data Figures 1-1, 1-2, 1-3, 1-4). In several cases, the coefficient for age dramatically differed from the mean of the distribution of individuals’ changes/slopes [i.e., (Brain_T2_-Brain_T1_)/(Age_T2_-Age_T1_)]. For example, had we controlled for ICV when analyzing cortical surface area, our analyses would have suggested that the surface area of the bilateral superior frontal, inferior temporal, and caudal middle frontal cortices decreased with age, even though the majority of individuals showed increases in surfaced area in these regions between both timepoints (Extended Data Figure 1-3). [Even though some individuals were removed from the longitudinal analyses due to their visit-specific data points being outliers (as described above), these models did include at least 95.2% of the individuals that contributed to the mean of these histograms (Extended Data Figures 1-1, 1-2, 1-3, 1-4).]

As a simple test to evaluate differences between the distributions of how individuals’ cortical metrics changed with age (i.e., individua slopes; Extended Data Figures 1-1, 1-2, 1-3, and 1-4) and the regression coefficients from each model (see Figure 1), we compared the signs (+/-) of the medians of the individual histograms to the signs (+/-) of the regression coefficients; intuitively, these signs should be matching, not opposing. However, in models that included the least optimal covariate, there were several instances of opposing signs [cortical thickness, controlling for mean cortical thickness: 20 of 68 brain regions (29.4%); cortical sulcal depth (ICV): 6 of 68 (8.8%); cortical surface area (ICV): 41 of 68 (60.3%) [and 15 of 68 (22.1%) when controlling for WBV]; and, cortical volume (total cortical volume): 29 of 68 (42.6%)]. In contrast, for each of the other covariate models across all four metrics, there was a maximum of 4 times (5.9%) in which the sign of the model’s regression coefficient for age did not match that of the distributional median of the individuals’ slopes.

**Figure 1.**
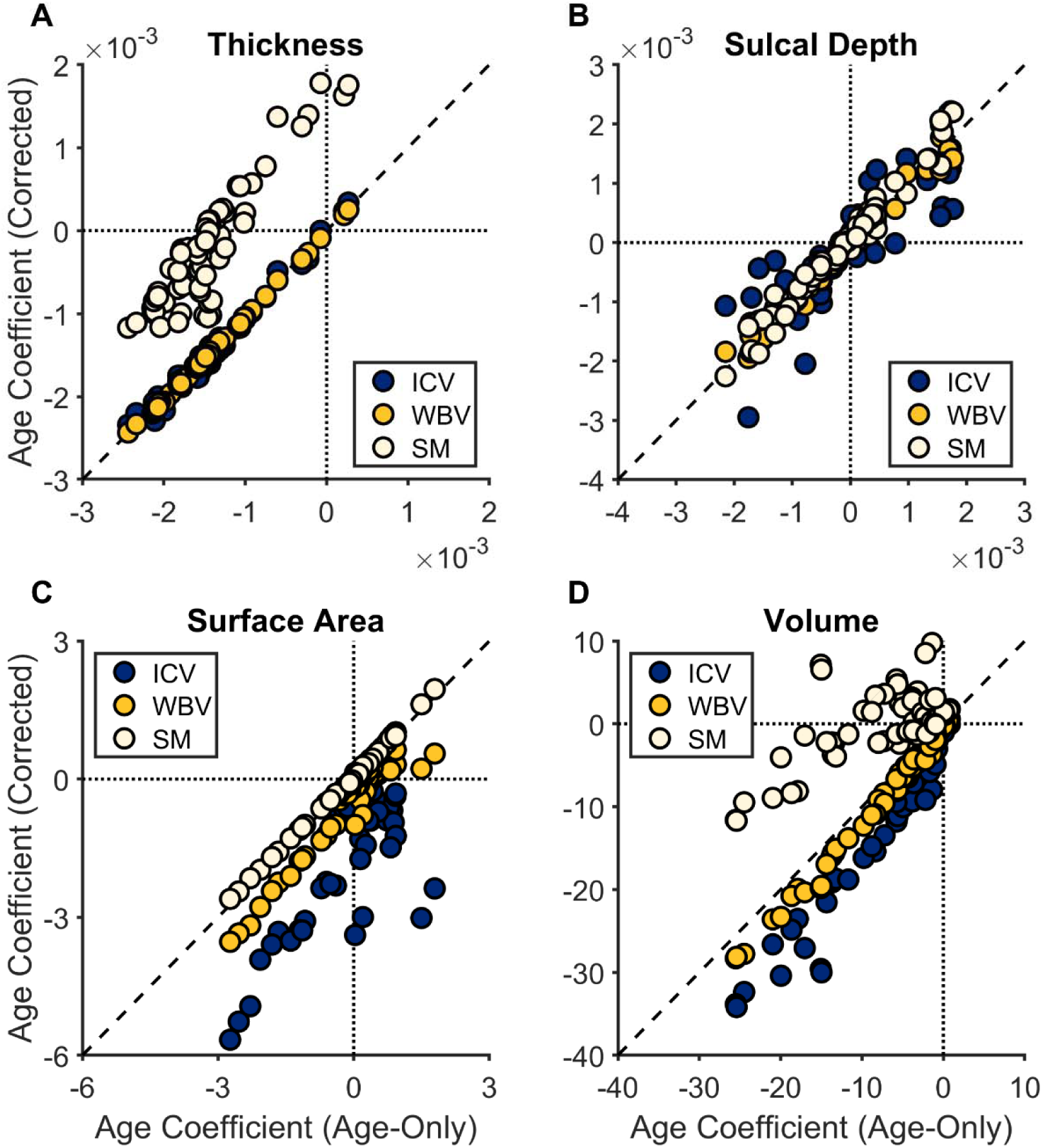
Comparison of associations between age and regional brain structure in linear mixed-effects models with (ordinate) and without (abscissa) whole-brain summary metrics. The abscissa shows the fixed-effects coefficient for “age” from the model *Brain Structure ∼ Age^2^ + (1|MRI Device) + (1|Participant)*; the ordinate shows the fixed-effects coefficient for “age” from the model *Brain Structure ∼ Age^2^ + Whole Brain Metric + (1|MRI Device) + (1|Participant)*, with whole-brain metric either being intracranial volume (ICV), whole brain volume (WBV), or an average/total summary metric (SM; i.e., mean thickness, meal sulcal depth, total surface area, total volume). The dashed vertical and horizontal lines partition the quadrants of the coordinate plane, and the dashed line is the unit diagonal.

At the population level, similar patterns were observed (Figure 1). For cortical thickness, the age coefficients from the WBV-corrected model were more highly correlated with the age coefficients from the age-only model (*r* > .99, *p* < .001) and more closely matched the corresponding values [total sum of squares (TSS) = 1.45 × 10^-7^] (ICV: *r* = .99, *p* < .001, TSS = 5.69 × 10^-7^; SM: *r* = 0.90, *p* < .001, TSS = 0.0001). The same was true for sulcal depth (WBV: *r* = .99, *p* < .001, TSS = 1.24 × 10^-6^; ICV: *r* = .86, *p* < .001, TSS = 1.60 × 10^-5^; SM: *r* = .99, *p* < .001, TSS = 2.53 × 10^-6^) and volume (WBV: *r* > .99, *p* < .001, TSS = 207.18; ICV: *r* = .97, *p* < .001, TSS = 1,766.23; SM: *r* = .64, *p* < .001, TSS = 4,672.22).

In contrast, SM-corrected models for surface area (i.e., correcting for total surface area) more closely matched the age-only models for surface area (SM: *r* > .99, *p* < .001, TSS = 0.26; WBV: *r* = .95, *p* < .001, TSS = 15.75; ICV: *r* = .72, *p* < .001, TSS = 168.42). Therefore, we controlled for WBV when modeling developmental trajectories of cortical thickness, sulcal depth, and volume, and we controlled for total area when modeling developmental trajectories of cortical surface area.

#### Associations between age and brain structure

Figure 2 shows the associations (i.e., effect sizes) between age and cortical structure by region, collapsed across sex-at-birth (Extended Data Figures 2-1, 2-2, 2-3, and 2-4 show the numerical values of these unstandardized regression coefficients and effect sizes). Here, there were highly significant changes in brain structure with age, with (1) global decreases in cortical thickness and volume (Figures 2A and 2D), (2) frontal-lobe increases and parieto-temporal decreases in sulcal depth (Figure 2B), and fronto-temporal increases, medial-occipital increases, and parieto-temporal decreases in surface area (Figure 2C).

**Figure 2.**
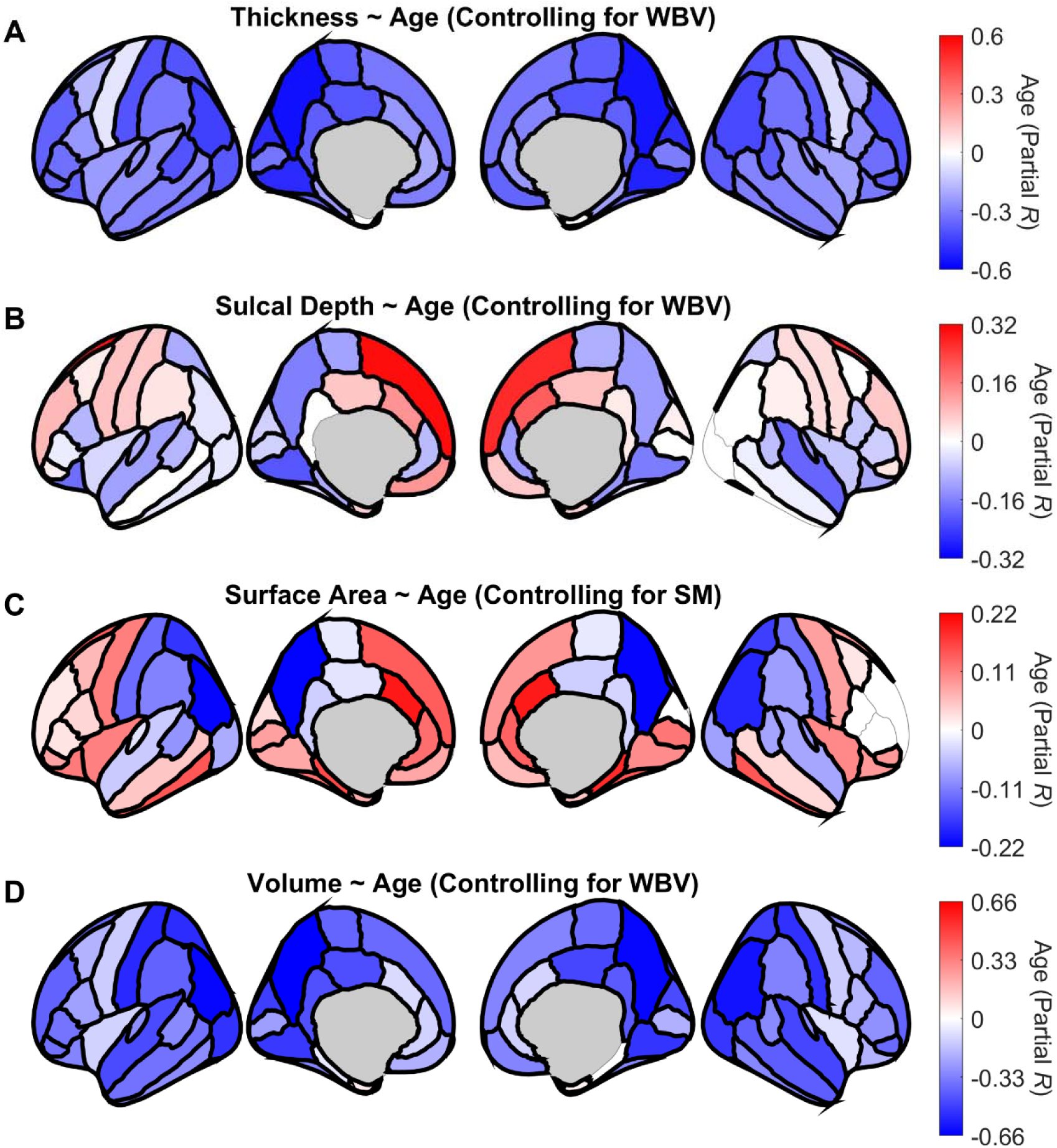
Associations between participant age and brain structure. Regions are color coded with respect to the color bars on the immediate right of each panel. (Note the different scales of the color bars.) Red- or blue-shaded regions indicate that those associations had *p*-values less than .05; regions that have thick borders indicate that those associations also passed false discovery rate (FDR) correction. These images were generated in MATLAB using data from the *ggseg* toolbox in R (Mowinckel & Vidal-Piñeiro, 2019).

#### Sex × Age interactions

Figures 3A-D show regional variation in how much sex-at-birth moderated associations between age and brain structure. In Figure 3, blue-shaded regions (or negative values on the color bar) reflect greater negative (or less positive) associations in individuals assigned female versus male sex-at-birth. In turn, red-shaded regions (or positive values on the color bar) reflect lesser negative (or greater positive) associations in individuals assigned male versus female sex-at-birth. (Extended Data Figures 3-1, 3-2, 3-3, and 3-4 show the numerical values of the unstandardized regression coefficients and effect sizes of the Sex × Age interaction terms.) While there were relatively minimal sex-at-birth-specific differences in developmental trajectories of sulcal depth and surface area (Figures 3B and 3C), youth assigned female sex-at-birth tended to have steeper declines in cortical thickness and volume with age relative to youth assigned male sex-at-birth (Figures 3A and 3D). With respect to the latter, the sex-at-birth-specific cortical-thickness trajectories were widespread, as were the cortical-volume trajectories, except for the lateral anterior parietal lobe.

**Figure 3.**
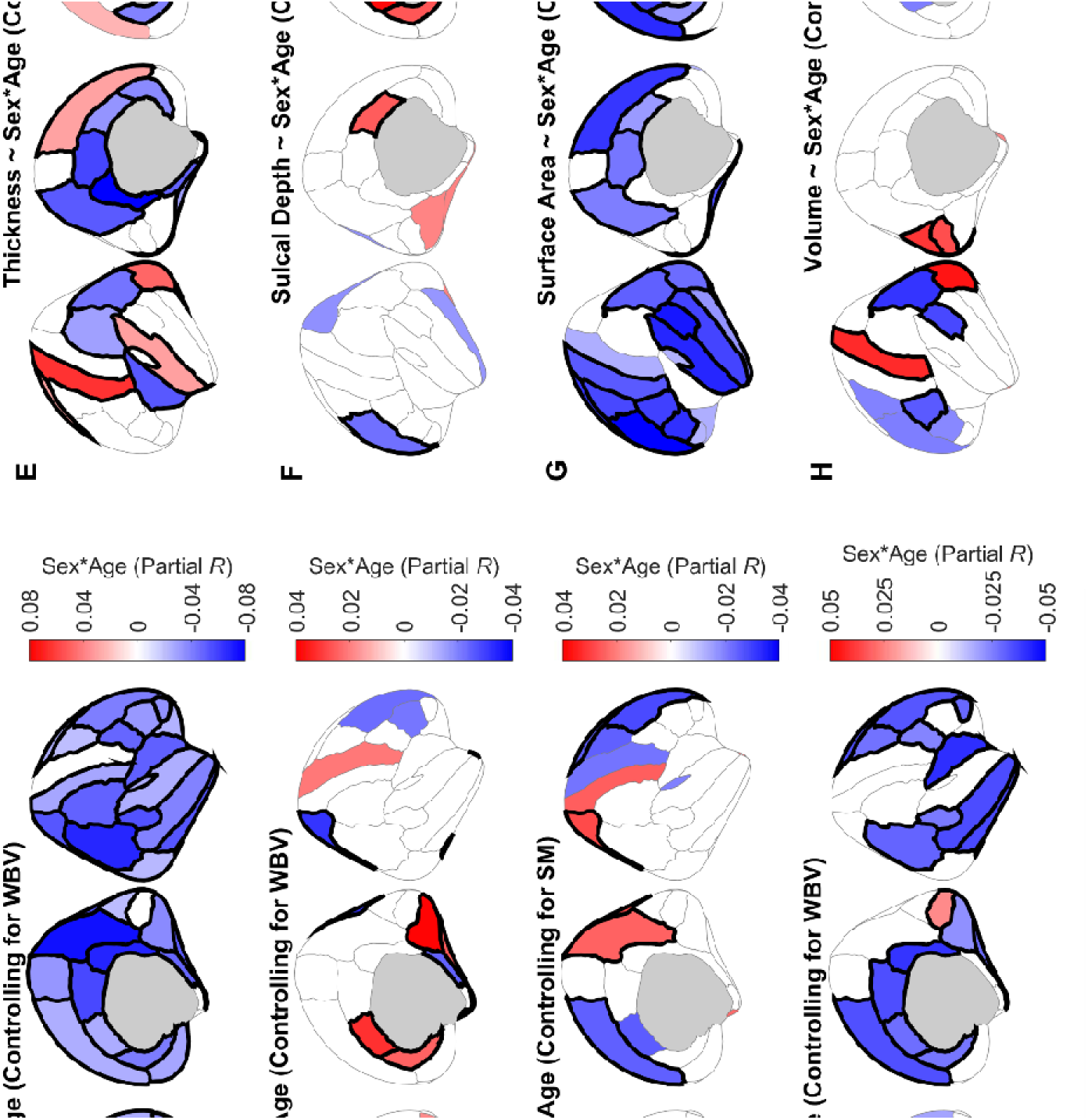
Sex × Age interactions on brain structure, controlling for different sample-specific whole-brain covariates. Panels A-D shows the interactions when controlling for the better-fitting sample-specific whole-brain covariates; Panels E-H, the worse-fitting sample-specific whole-brain covariates. Regions are color coded with respect to the color bars on the immediate right of each panel. (Note the different scales of the color bars.) Red- or blue-shaded regions indicate that those associations had *p*-values less than .05; regions that have thick borders indicate that those associations also passed FDR correction. Blue-shaded regions reflect greater negative (or lesser positive) associations in individuals assigned female sex at birth. Red-shaded regions reflect greater positive (or lesser negative) associations in individuals assigned male sex at birth. These images were generated in MATLAB using data from the *ggseg* toolbox in R (Mowinckel & Vidal-Piñeiro, 2019).

In contrast, Figures 3E-H show the distributions and strengths of Sex × Age interactions on cortical structure had we controlled for the whole-brain covariates linked to coefficients that were most different from the age-only coefficients (i.e., mean cortical thickness, total cortical volume, and ICV for sulcal depth and surface area; Figure 1). While patterns pertaining to sulcal depth were relatively consistent, there were stark differences for thickness, surface area, and volume. For example, while there remained Sex × Age interactions on cortical thickness, these results were more varying in direction and less widespread (Figure 3E). Instead, this analysis showed more widespread and consistent Sex × Age interactions on surface area, which, in conjunction with Extended Data Figure 1-3, would have suggested (potentially erroneously) that individuals assigned female sex-at-birth showed even greater declines in surface area with age compared to those assigned male sex-at-birth.

### Annual Percentage Change (APC)

Given distinct developmental trajectories of cortical thickness and surface area, coupled with how much these patterns depend on covariate selection (Figure 3, Extended Data Figures 1-1, 1-2, 1-3, 1-4), we conducted a second, more holistic/omnibus set of analyses to measure how the cortex is changing relative to itself. Specifically, we evaluated (1) whether there were sex-at-birth-specific differences in how rates of cortical thickening/thinning (thickness) compared to rates of cortical expansion/contraction (surface area) in each region, (2) whether there were omnibus sex-at-birth-specific within-region relationships between these two processes, and (3) how within-region thickness/surface-area trajectories varied by sex-at-birth and age.

Figures 4-5 show relationships between the regional APCs of cortical thickness and surface area in each of 6,323 participants, 5,362 of whom were scanned at both timepoints on the same MRI scanner (84.8%). There were 410,538 data points for surface area (outlier or missing *n* = 19,426) and 411,534 for cortical thickness (outlier or missing *n* = 18,430). In both hemispheres for both groups, APC_CT_ was highly (and similarly) correlated with the APC_CSA_ (Male/Left: *r* = -0.19; Male/Right: *r* = -0.20; Female/Left: *r* = -0.19; Female/Right: *r* = -0.19; all *p*’s < .001). These patterns were similar for individuals scanned within the same machine (Male/Left: *r* = -0.19; Male/Right: *r* = -0.20; Female/Left: *r* = -0.18; Female/Right: *r* = -0.18; all *p*’s < .001). For individuals assigned male sex-at-birth (and those scanned on the same machine), 39.3% (39.3%) of the data points (across both hemispheres) were in the cortical thinning/expanding quadrant of the coordinate plane, followed by 30.5% (30.8%) in the thinning/contracting quadrant, 15.5% (15.2%) in thickening/contracting quadrant and 13.4% (13.4%) in the thickening/expanding quadrant. For individuals assigned female sex-at-birth, 37.6% (37.7%) of the data points were in the cortical thinning/expanding quadrant, followed by 35.4% (35.9%) in the thinning/contracting quadrant, 14.6% (14.2%) in the thickening/contracting quadrant, and 11.1% (10.9%) in the thickening/expanding quadrant. Thus, distributionally, there were similar whole-brain patterns of individual-differences results by sex-at-birth.

**Figure 4.**
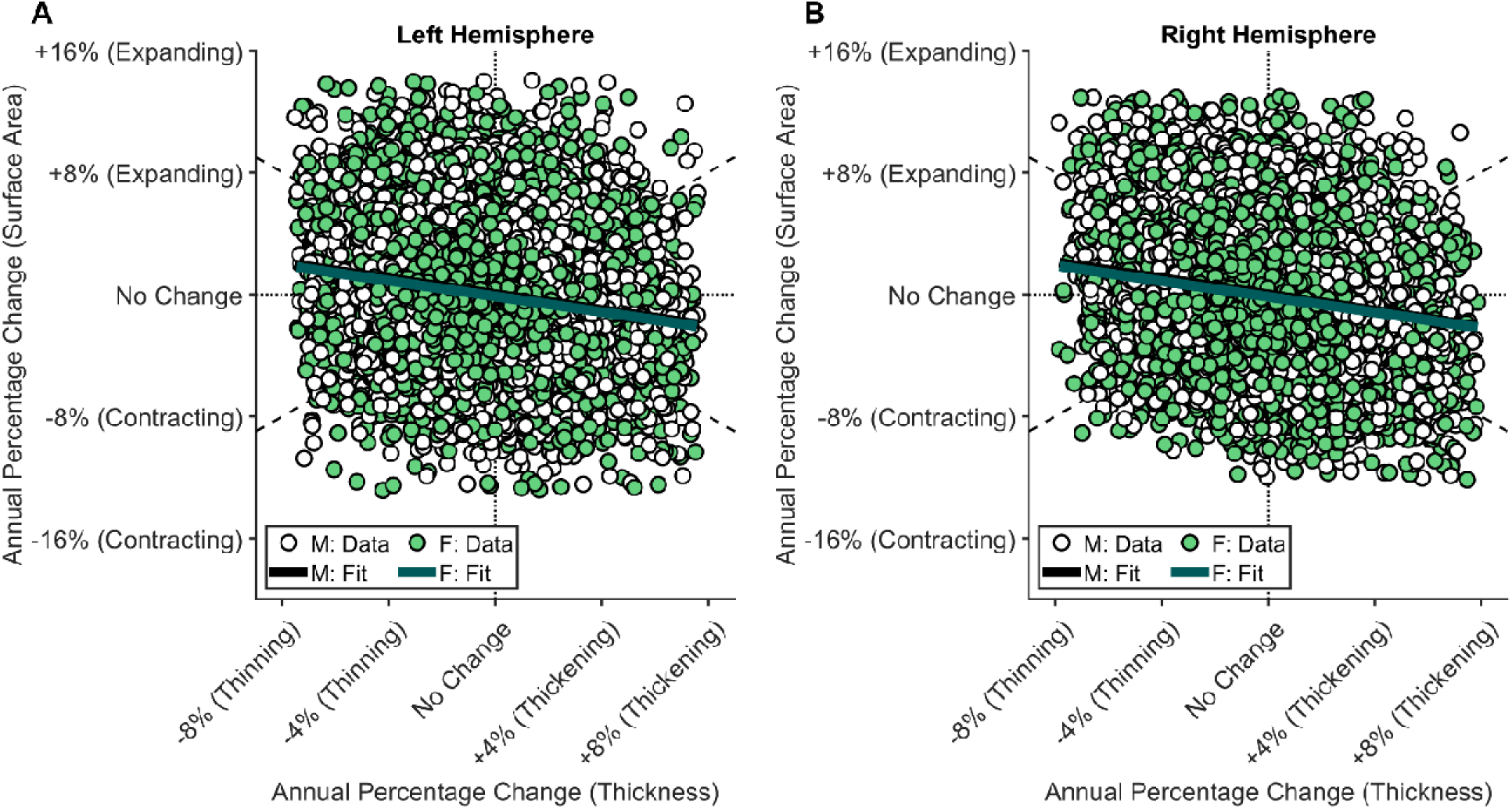
Individual differences in regional annual percentage change in cortical thickness and surface area. Each data point represents one region from one individual, color coded by sex-at-birth and split into the left hemisphere (**A**) and right hemisphere (**B**). The thick, solid lines in each graph are the best fit linear fits per simple regression (Surface Area ∼ 1+ Thickness). The dotted horizontal and vertical lines partition the quadrants of the coordinate plane, and the dashed lines are the unit diagonals.

**Figure 5.**
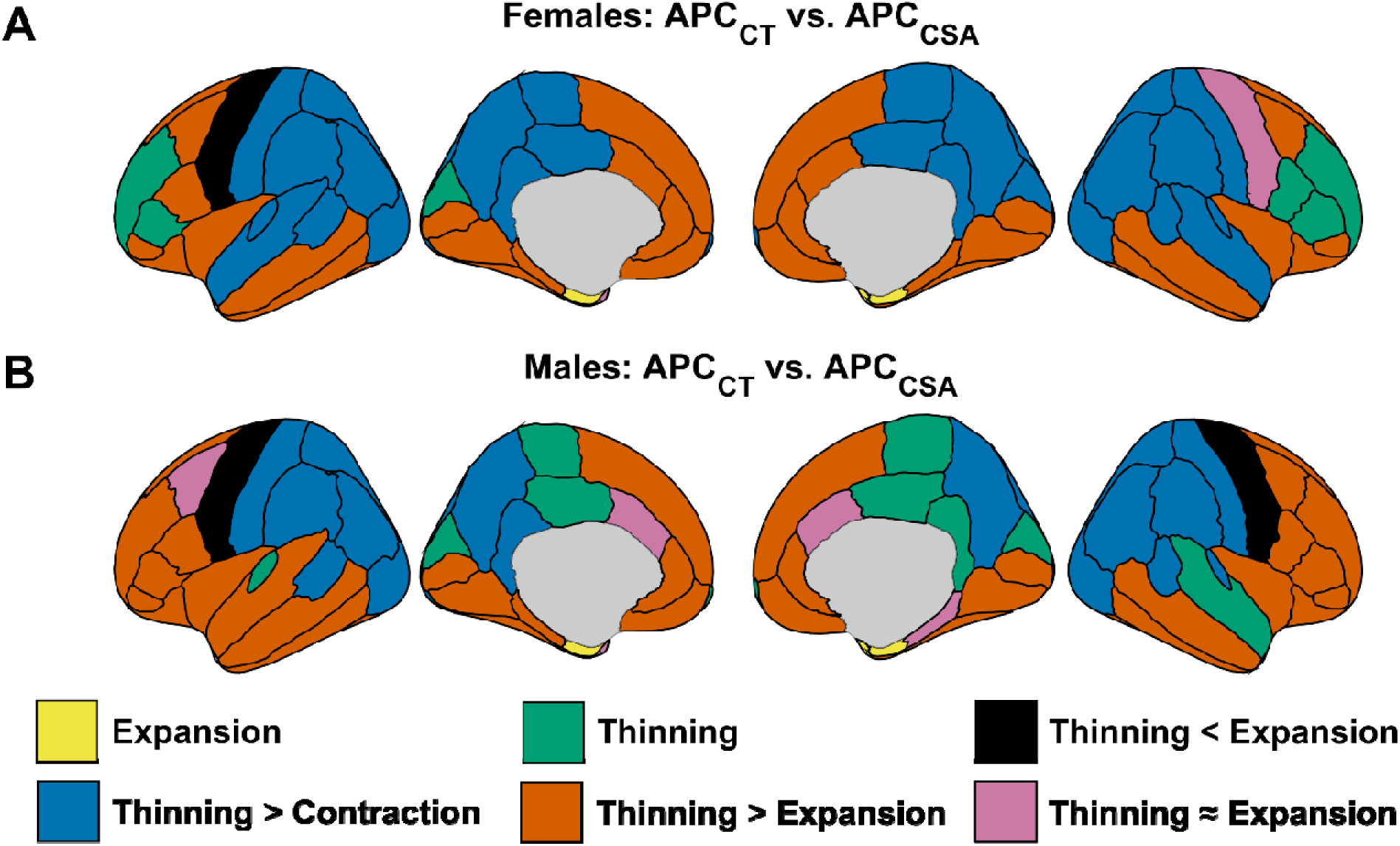
Regional differences in rates of developmental change in cortical thickness and surface area per annual percentage change. Regions are color coded with respect to differential rates of cortical thinning/thickening vs. cortical contraction/expansion, based on whether the annual percentage change rates were significantly different from 0 and whether the rate of thinning/thickening was significantly different than that of contraction/expansion. Regions that were color coded for only one type of change (i.e., “Expansion”, “Thinning”) indicates that the opponent process’s annual percentage change was not significantly different from 0.

### Comparing APC_CT_ to APC_CSA_

Regionally, the brains of individuals assigned male sex-at-birth changed in a generally similar fashion to those assigned female sex-at-birth (Figure 5). For example, cortical thinning outpaced (1) surface expansion in several fronto-temporal regions and (2) surface contraction in parietal and lateral temporal regions. With some exception, there was minimal cortical asymmetry in the collective patterning of APC_CT_ versus APC_CSA_. Further, the most notable categorical sex-at-birth-specific difference in these designations was in the left superior temporal cortex, in which, in individuals assigned female sex-at-birth, cortical thinning outpaced cortical contraction, while in individuals assigned male sex-at-birth, the rate of cortical thinning exceeded that of cortical expansion.

Interestingly, while the anterior frontal cortex of individuals assigned male sex-at-birth exhibited significant thinning outpacing significant expansion, that of individuals assigned female sex-at-birth exhibited significant thinning. Similarly, in the posterior frontal cortex, in individuals assigned female sex-at-birth, significant thinning outpaced significant contraction, while individuals assigned male sex-at-birth primarily showed significant thinning. With respect to categorical representativeness, aside from the bilateral frontal pole in individuals assigned female sex-at-birth, the proportion of individuals (by region and sex-at-birth) whose trajectories matched the group-level patterns (Figure 5) exceeded random chance (Extended Data Figure 5-1), suggesting that the group-level patterns were at least partially representative of its individuals. These patterns were similar when considering only the individuals scanned by the same machine at both timepoints (Extended Data Figures 5-2 and 5-3).

### Circular analyses: APC_CT_-APC_CSA_ trajectories by age

Thus far, these results offer (at least) two (not necessarily mutually exclusive) interpretations: (1) there are vast sex-at-birth-specific differences in how cortical thickness is changing with age at this stage of adolescence (Figure 3), even though (2) the extent of these differences may be diminished by the degree of overlap in such differences (Figures 4-5). In other words, differences in developmental change conveyed by singular cortical metrics may not necessarily mean that the brains of individuals assigned female or male sex-at-birth are changing in categorically different ways; instead, there may be differences in the timing of these changes, which can be reconciled via further waves of longitudinal study. Therefore, rather than considering whether specific cortical metrics are changing at certain rates (and whether those particular metrics are happening at significantly greater or lesser rates in one group than another), we analyzed co-directional distributions of developmental change in cortical thickness (APC_CT_) and surface area (APC_CSA_) by region, hemisphere, and sex-at-birth (see Figure 4).

Collapsed across all regions per hemisphere, there were relatively similar distributions by sex-at-birth across octants (Figure 6), in which cortical thinning was paralleled by both cortical expansion and contraction (with a slight bias toward cortical expansion in individuals assigned male sex-at-birth and cortical contraction in those assigned female sex-at-birth). Regionally, while there were Benjamini-Hochberg-corrected main effects of sex-at-birth for 55 of the 68 regions (Extended Data Figure 7-1), the corresponding distributions were relatively similar (Figure 7), with sex-at-birth accounting for a maximum of 1.04% of the variance in the APC_CT_- APC_CSA_ circular distributions.

**Figure 6.**
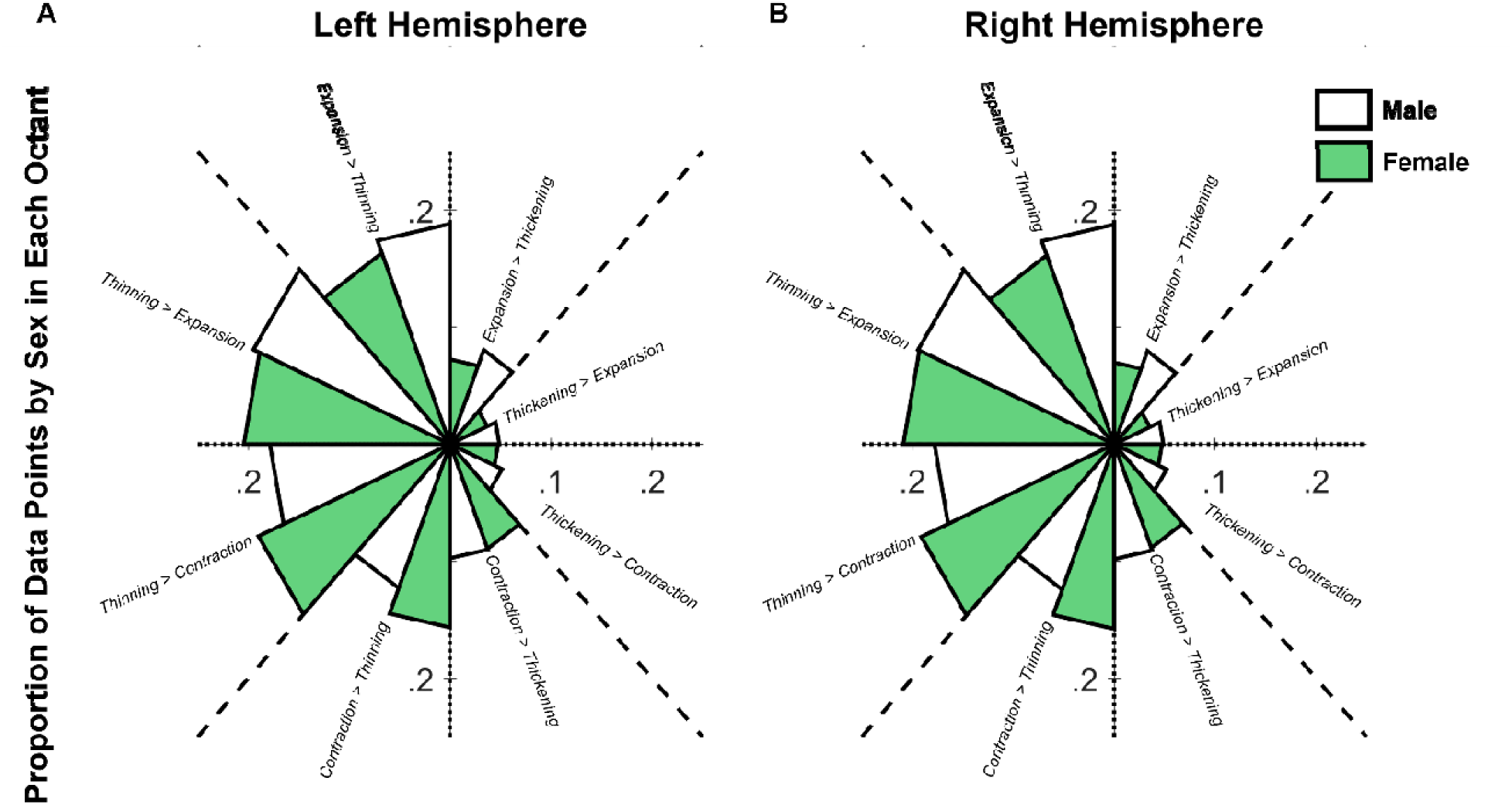
Distribution of annual percentage change in cortical thickness and surface area. Each segment represents the proportion of individuals by sex-at-birth within each octant, color coded by sex-at-birth and split into the left hemisphere (**A**) and right hemisphere (**B**). The dotted horizontal and vertical lines partition the quadrants of the coordinate plane, and the dashed lines are the unit diagonals.

**Figure 7.**
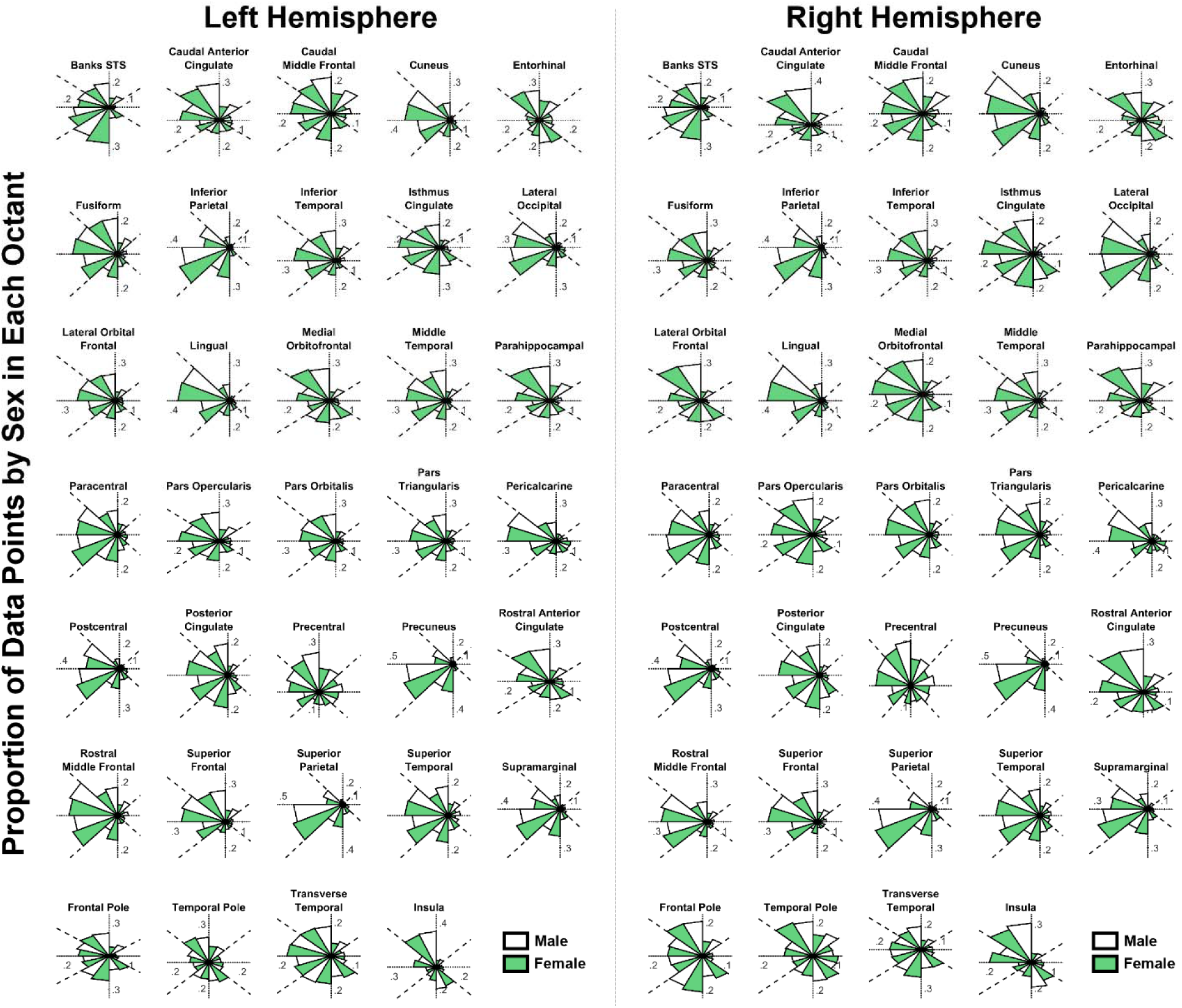
Regional distributions of annual percentage change in cortical thickness and surface area. Each segment represents the proportion of individuals by sex-at-birth within each octant, color coded by sex-at-birth and split into the left and right hemispheres. The dotted horizontal and vertical lines partition the quadrants of the coordinate plane, and the dashed lines are the unit diagonals.

Interestingly, visual inspection of the global and regional distributions in Figures 6-7 suggest an intriguing difference by sex-at-birth, particularly considering the minimal variance accounted for by sex-at-birth. Specifically, rather than there being markedly different APC_CT_- APC_CSA_ distributions by sex-at-birth, the data for those assigned female sex-at-birth simply appear to be rotated counterclockwise relative to those assigned male sex-at-birth. Accordingly, when the data in Figure 7 were further split by age and sex-at-birth (Figure 8), it was apparent that, generally, both sexes showed similar trends with age, in which cortical thinning and expansion at younger adolescent ages transitioned into cortical thinning and contraction at older adolescent ages (i.e., downward slopes in Figure 8). However, in several regions, this transition seemed to occur at slightly older ages in those assigned male sex-at-birth. For example, in the bilateral rostral middle frontal and superior temporal cortices, the thinning/expansion-to-thinning/contraction transition begins slightly later in adolescence in those assigned male versus female sex-at-birth.

**Figure 8.**
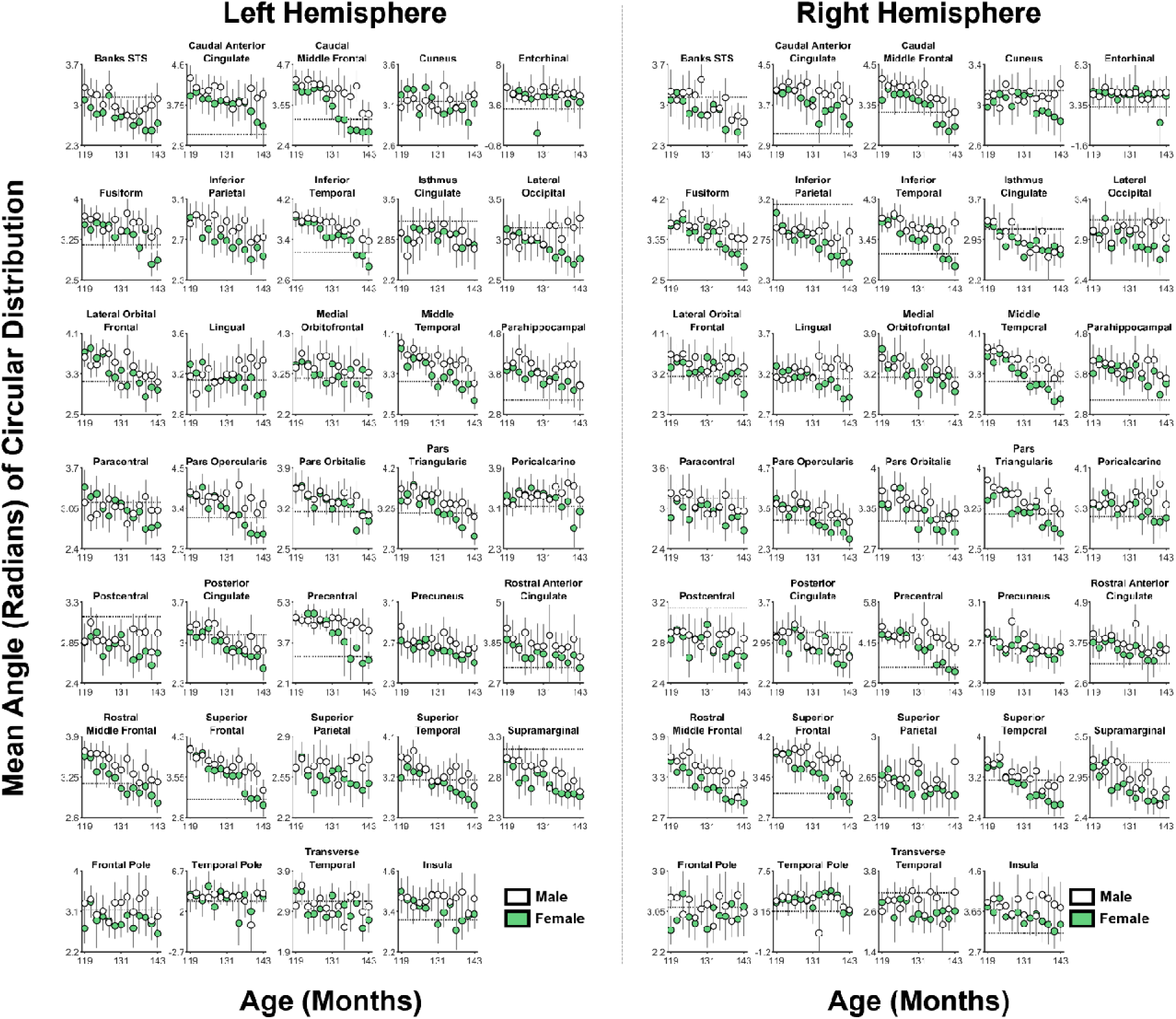
Mean angles (directions) of annual percentage change in cortical thickness and surface area by sex-at-birth and age (in months). Intuitively, each data point represents the mean direction ± 95% confidence intervals of the data in Figure 7, had the data in Figure 7 also been partitioned by participant age. Age (in months) for each participant was the midway point between each of their ages at the baseline and two-year follow-up appointments. The dashed horizontal line (*y* = π) represents cortical thinning with no corresponding change in cortical surface area. For the bilateral temporal pole and entorhinal cortices, the values less than 0 and/or greater than 2π are not erroneous but artifacts of converting data points from a circular to non-circular scale.

## Discussion

We have reported sophisticated yet integrated analyses of cortical maturation, specifically considering the individuals that comprise population-level changes. Importantly, we have shown that covariate selection matters when interpreting developmental cortical maturational trajectories. Moreover, we have provided a more nuanced approach to measuring developmental trajectories by more aggregately evaluating multivariate cortical change. Overall, while there was substantial cortical development (Figures 2-3), a more interesting pattern of results emerged from the APC_CT_-APC_CSA_ bivariate analyses: Sex-at-birth differences in cortical development may be more so one of “when” then “how much”.

First, we evaluated how sex-at-birth moderated cortical change in adolescence, in conjunction with past research on multivariate adolescent cortical development (e.g., Blakemore, 2012; Brown et al., 2012; Mills et al., 2021; Raznahan et al., 2011; Schnack et al., 2015; Tamnes et al., 2017; Vijayakumar et al., 2016). Generally, cortical thinning happened at faster rates in those assigned female than male sex-at-birth, with less pervasive uniformity in how sex-at-birth moderated sulcal-depth and surface-area development (Figures 3A-3D). Given frequently reported differences in brain size by sex (e.g., Eliot et al., 2021) and decisions to statistically control for these differences (see, e.g., Mills et al., 2016), we tested how these associations depended on which covariate was included in the model, premised on the reasonable assumption that fixed-effects coefficients should generally capture what is occurring at the individual level; otherwise, group-level coefficients fail to summarize the individuals comprising that group. Here, WBV was the more optimal covariate selection for change in cortical thickness, sulcal depth, and volume; total cortical surface area was more optimal for regional surface-area change (Figure 1). Importantly, a different selection would have altered interpretations of both age-related associations and Sex×Age interactions (Figure 3), with the former being most noteworthy for surface area: Controlling for ICV would have revealed cortical change in the opposite direction to that shown by the majority of participants (Extended Data Figure 1-3), a neuroanatomical example of Simpson’s paradox (Simpson, 1951).

Previous reports have emphasized how theory and hypothesis should inform covariate selection in neuroimaging analyses (O’Brien et al., 2011; O’Brien et al., 2006), but our results underscore the importance of statistically rationalizing covariate selection based on patterns in each dataset. To our knowledge, such rigorous yet simple testing is underutilized in developmental neuroscience. One oft-cited report compared how controlling for ICV or WBV (as covariates or by proportion) altered interpretation of developmental maturation of cortical volume (Mills et al., 2016). Similarly, here, different decisions may have led to different conclusions [e.g., a blanket choice to control for summary metrics (i.e., mean thickness, total surface area, total volume) would have promoted conclusion that there is little directional or regional uniformity in how the adolescent brain is changing; Figures 3C, 3E, 3H]. Thus, expanding on previous arguments (see Barnes et al., 2010; Mills et al., 2016; O’Brien et al., 2011; O’Brien et al., 2006), these decisions should not solely rely on expectation, population comparison, and inference, but also on how well our models are most effectively capturing individual data (Extended Data Figures 1-1, 1-2, 1-3, 1-4). [While a comprehensive review of covariate selection/comparison across the developmental-neuroscience literature extends beyond our scope, we cursorily reviewed 861 ABCD-related publications identified by the study’s primary website (https://abcdstudy.org/publications, as of 25 January 2024), 78 of which we viewed as conducting analyses methodologically pertinent to ours (e.g., cortical ROIs across the whole brain, etc.) (Table 3). The relevant covariates for body/brain size (or lack thereof) largely varied, with only a few publications exploring different covariates. For example, using baseline ABCD data, Dhamala et al. (2022) showed that ICV-corrected thickness models and ICV-uncorrected surface-area and volume models outperformed ICV-*uncorrected* thickness models and ICV-*corrected* surface-area and volume models, respectively, partially congruent with how ICV was associated with model differences here (Figure 1).]

**Table 3.** ABCD publications with whole-brain cortical region-of-interest analyses relevant to those conducted here (as of 25 January 2024).

|  | Thickness<br>(T) | Sulcal<br>Depth (SD) | Surface Area<br>(SA) | Volume (V) |
| --- | --- | --- | --- | --- |
| Adise, Marshall, Hahn, et al. (2023) | ICV | -- | ICV | -- |
| Adise, Marshall, Kan, et al. (2023) | NR | -- | -- | -- |
| Ahmed, Cano, Sánchez, Hu, Gonzalez, et al. (2023) | NR | -- | -- | -- |
| Ahmed, Cano, Sánchez, Hu and Ibañez (2023) | NR | -- | -- | -- |
| Alnæs et al. (2020) | NR | NR | NR | ICV |
| Blok et al. (2023) | -- | -- | -- | ICV (±) |
| Bohon and Welch (2021) | -- | -- | -- | ICV (±), TBV (±) |
| Brooks et al. (2023) | NR | -- | -- | NR |
| Bustamante et al. (2022) | -- | -- | -- | Total V |
| Cheng et al. (2021) | NR | -- | ICV (±), Total SA (±) | ICV (±), Total V (±) |
| Cserbik et al. (2020) | NR | -- | NR | -- |
| Cui et al. (2023) | ICV | -- | ICV | ICV |
| Dai et al. (2022) | -- | -- | NR | NR |
| Dhamala et al. (2022) | ICV (±) | -- | ICV (±) | ICV (±) |
| Durham et al. (2023) | -- | -- | -- | ICV (±) |
| Durham et al. (2021) | -- | -- | -- | ICV (±), Total V (±) |
| Fan et al. (2023) | ICV, Mean T | ICV | ICV | ICV |
| Fernandez-Cabello et al. (2022) | NR | -- | NR | -- |
| Gaus et al. (2023) | Supratent. V | -- | -- | Supratent. V |
| Ge et al. (2023) | NR | -- | ICV | -- |
| Hackman et al. (2021) | -- | -- | Total SA (±) | ICV |
| Hatoum et al. (2021) | Mean T | -- | Mean SA | Mean V |
| He et al. (2023) | Mean T | -- | Total SA | Total V |
| Hernandez et al. (2022) | Mean T (±) | -- | Mean SA (±) | -- |
| Isaiah et al. (2021) | NR | -- | NR | NR |
| Isaiah et al. (2023) | -- | -- | -- | NR |
| Jeong et al. (2021) | Mean T | -- | -- | ICV (±), Total V (±) |
| Jeong et al. (2022) | Mean T (±) | -- | -- | ICV (±) |
| Ji et al. (2024) | ICV | -- | -- | ICV |
| Kaltenhauser et al. (2023) | ICV | -- | -- | -- |
| Karcher et al. (2023) | ICV | -- | -- | -- |
| Karcher et al. (2022) | ICV | -- | ICV | ICV |
| Karcher et al. (2021) | ICV | -- | ICV | ICV |
| Kuang et al. (2023) | ICV, Mean T | -- | ICV, Total SA | -- |
| Kulisch et al. (2023) | Mean T | -- | Total SA | Total V |
| Lahey et al. (2024) | -- | -- | -- | ICV (±) |
| Laurent et al. (2020) | ICV | -- | -- | -- |
| Lees et al. (2020) | NR | -- | NR | ICV (±) |
| Lees et al. (2021) | NR | -- | ICV | ICV |
| Li et al. (2024) | -- | -- | -- | NR |
| Luo et al. (2023) | -- | -- | -- | ICV |
| Luo et al. (2022) | -- | -- | -- | ICV |
| Ma et al. (2023) | *NR* | -- | -- | -- |
| Ma et al. (2022) | -- | -- | -- | *NR* |
| MacSweeney et al. (2023) | WBV (±) | WBV (±) | WBV (±) | WBV (±) |
| Makowski et al. (2022) | Mean T (±) | -- | Total SA (±) | -- |
| Menken et al. (2023) | ICV | -- | ICV | ICV |
| Mewton et al. (2023) | -- | -- | -- | *NR* |
| Mewton et al. (2022) | Mean T (±) | -- | Mean SA (±) | Total V (±) |
| Moreau et al. (2022) | Mean T (±) | -- | ICV | -- |
| Morys et al. (2023) | *NR* | -- | -- | ICV |
| Nath et al. (2023) | Hemi. Mean T<br>(±) | -- | Hemi. Total<br>SA (±) | -- |
| Pagliaccio et al. (2021) | Height* | -- | -- | -- |
| Petrican et al. (2023) | *NR* | -- | -- | -- |
| Rakesh et al. (2022) | Mean T (±) | -- | TBV (±) | -- |
| Rakesh et al. (2023) | *NR* | -- | TBV | -- |
| Ronan et al. (2020) | Brain vol. | -- | -- | -- |
| Rutherford et al. (2022) | *NR* | -- | -- | -- |
| Shadrin et al. (2021) | Mean T | -- | Total SA | -- |
| Shen et al. (2023) | -- | -- | *NR* | -- |
| Shen et al. (2021) | ICV | ICV | ICV | ICV |
| Sun et al. (2024) | ICV (±) | -- | ICV (±) | ICV (±) |
| Teeuw et al. (2023) | -- | -- | -- | TBV (±) |
| Tomasi and Volkow (2021) | ICV | -- | -- | ICV |
| Vaughn et al. (2021) | *NR* | -- | -- | -- |
| Vidal-Ribas et al. (2021) | Mean T | -- | -- | -- |
| Wang et al. (2020) | *NR* | -- | *NR* | *NR* |
| Wang et al. (2023) | *NR* | -- | *NR* | *NR* |
| Wen, Qu, et al. (2023) | -- | -- | -- | ICV |
| Wen, Shu, et al. (2023) | ICV | -- | ICV | ICV |
| Westwater et al. (2023) | ICV | -- | ICV | -- |
| Wiglesworth et al. (2023) | *NR* | -- | -- | -- |
| Wu et al. (2022) | -- | -- | -- | ICV |
| Yang et al. (2023) | -- | -- | -- | ICV |
| Yang et al. (2022) | *NR* | -- | *NR* | *NR* |
| Yu et al. (2023) | ICV | -- | -- | -- |
| Zhang et al. (2022) | *NR* | *NR* | *NR* | -- |
| Zhao et al. (2022) | Total T (±) | -- | *NR* | -- |

*Note.* \* = While analyses in other publications included body-mass index as a covariate, this publication specifically noted that height was used as a covariate to control for body size. Hemi. = Hemispheric. ICV = Intracranial volume. *NR* = Not reported. Supratent. = Supratentorial. TBV = Total brain volume. Vol. = Volume. WBV = Whole brain volume. ± = Evidence for similar analyses being conducted with and without that covariate.

The interpretational fallout of covariate selection speaks to whether past and presently reported maturational cortical differences are real or if they are simply artifacts of the chosen covariate. In the vein of sensitivity analyses, one solution would be to identify regions that maintained significant effects in the same direction regardless of covariate. Indeed, in doing so here, with reference to WBV models for thickness, sulcal depth, and volume, and SM models for surface area, such robustness of significant effects is present in 18 (of 68) ROIs for cortical thickness, 14 ROIs for sulcal depth, 4 ROIs for surface area, and 7 ROIs for cortical volume. Therefore, while cortical developmental trajectories may be influenced by sex-at-birth, such differences may be less widespread, absolute, large, or small than any one analysis indicates (cf. DeCasien et al., 2022; Eliot et al., 2021; Williams et al., 2021). However, as discussed below, it may not be that such sex-at-birth differences are nonexistent, but that the critical difference is when in adolescence those changes start to happen (i.e., beyond the ages of all participants with ABCD 4.0 data).

Ultimately, an approach to circumvent dilemmas of choosing between distinct yet interrelated covariates would be to treat each individual as their own control (i.e., how individuals are changing relative to themselves). Fortunately, ABCD’s longitudinal design facilitates this option, as best exemplified by the multimodal analyses conducted by Bottenhorn et al. (2023), in which their reported variabilities of individual differences in microstructural, macrostructural, and functional MRI metrics reflects the value and importance of evaluating such idiosyncrasies. Accordingly, in our second set of analyses, we computed and compared APC_CT_ and APC_CSA_ to evaluate sex-at-birth differences in these bivariate distributions. In a non-ABCD sample, Mills et al. (2021) reported sex differences in developmental trajectories of global cortical metrics, as well as in individual-differences relationships between global brain metrics (e.g., total gray matter volume) and how much that global brain metric changed with age. Interestingly, while Wierenga et al. (2019) reported several regional sex-by-age interactions on cortical thickness and surface area, the vast majority of those corresponding best-fitting models for cortical structure did not include these interactions, paralleling their primary findings of greater sex differences in the variance (not means) of brain structure.

Here, there were statistically significant effects of sex-at-birth, but the omnibus group APC_CT_-APC_CSA_ distributional patterns were generally comparable (Figures 5-7). Notably, the corresponding regions with the largest effects of sex-at-birth were in the frontal and temporal cortices (Extended Data Figure 7-1), which generally corresponded to the sex-at-birth differences in categorization of cortical thinning/thickening versus contraction/expansion (Figure 5). For instance, while those assigned female sex-at-birth tended to only show significant thinning in the bilateral rostral middle frontal and pars triangularis cortices, those assigned male sex-at-birth showed both significant thinning and expansion, with the former outpacing the latter. As implied by the data in Figure 8, strictly cross-sectional analyses at any given age (particularly older adolescent ages) would likely have revealed large sex-at-birth differences in cortical development (i.e., differences on the vertical axis, as essentially shown in Figure 3). However, in consideration of the overall patterning of these data (Figure 8), it seems that instances of sex-at-birth differences in cortical development may actually be one of “when” rather than “how much”. In other words, the key sex-at-birth difference in adolescent cortical development may be *when* that cortical development starts as opposed to *how much* cortical development ultimately happens during any snapshot of time. Indeed, Lenroot et al. (2007) reported that total cerebral volume peaked ∼4 years earlier in female than male participants, and Koolschijn and Crone (2013) suggested that surface expansion may be completed earlier in development in female than male individuals, potentially due to age-related differences in pubertal stage (Beck et al., 2023). Thus, nominal categorization of cortical change (Figure 5) may be less reflective of different developmental trajectories but simply different stages of relatively similar trajectories, with frontal and temporal lobar surface expansion ending earlier in those assigned female than male sex-at-birth. As similarly conveyed by Beck et al. (2023), while we are limited in only having two datapoints per participant in our analyses, the continued collection and release of ABCD data will permit, in our context, opportunity to potentially “align” two-year developmental segments of cortical change to elucidate these possibilities.

In conclusion, while previous research has reported relationships between developmental trajectories of cortical thickness, surface area, and volume in children and adults (Lyall et al., 2015; Storsve et al., 2014; Tamnes et al., 2017), our results provide foundation for reconceptualization and reconsideration of how to interpret univariate cortical maturation. Reconsidering one of ABCD’s key objectives (“develop national standards for normal brain development in youth, by defining the range and pattern of variability in trajectories of brain development”) (Jernigan, Brown, & ABCD Consortium Coordinators, 2018, p. 2), while there is reason to consider analyses of cortical thickness and surface area separately (Panizzon et al., 2009), much greater insight may be gained through, e.g., vector analyses that not only reflect singular metrices but the relationship between multivariate cortical trajectories. As ABCD progresses, there will be greater opportunities to more precisely model individual trajectories of brain, cognitive, physiological, social, emotional, and behavioral development in future research.

## Acknowledgments

Data used in the preparation of this article were obtained from the Adolescent Brain Cognitive Development^SM^ (ABCD) Study (https://abcdstudy.org), held in the NIMH Data Archive (NDA). This is a multisite, longitudinal study designed to recruit more than 10,000 children age 9-10 and follow them over 10 years into early adulthood. The ABCD Study® is supported by the National Institutes of Health and additional federal partners under award numbers U01DA041048, U01DA050989, U01DA051016, U01DA041022, U01DA051018, U01DA051037, U01DA050987, U01DA041174, U01DA041106, U01DA041117, U01DA041028, U01DA041134, U01DA050988, U01DA051039, U01DA041156, U01DA041025, U01DA041120, U01DA051038, U01DA041148, U01DA041093, U01DA041089, U24DA041123, U24DA041147. A full list of supporters is available at https://abcdstudy.org/federal-partners.html. A listing of participating sites and a complete listing of the study investigators can be found at https://abcdstudy.org/consortium_members/. ABCD consortium investigators designed and implemented the study and/or provided data but did not necessarily participate in the analysis or writing of this report. This manuscript reflects the views of the authors and may not reflect the opinions or views of the NIH or ABCD consortium investigators. The ABCD data repository grows and changes over time. The ABCD data used in this report came from 10.15154/1523041. We would like to specifically thank Dr. Megan Herting (USC) for her comments on earlier drafts of the manuscript.

