## Extended Data for "Longitudinal sex-at-birth and age analyses of cortical structure in the ABCD Study^®^"

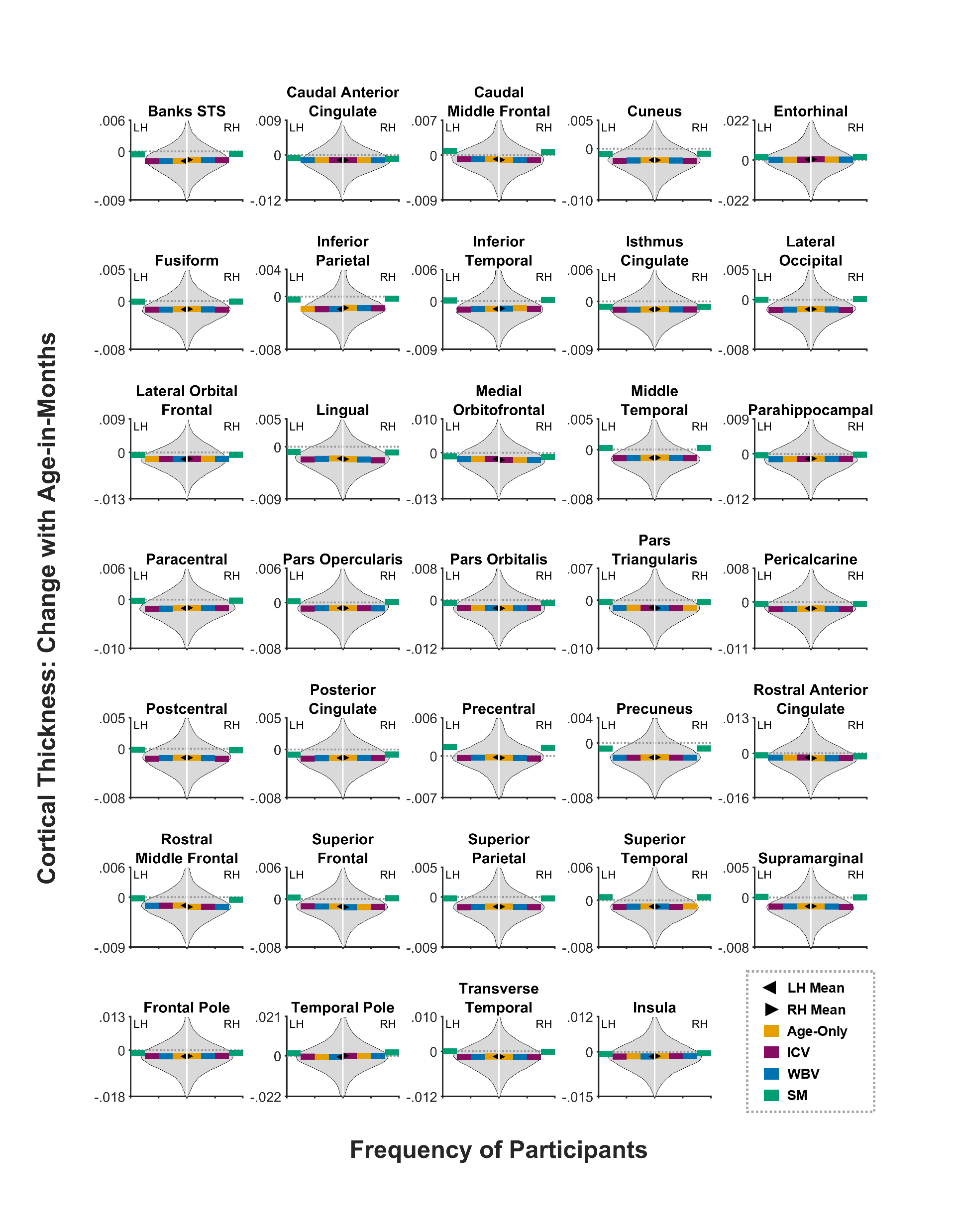

**Extended Data Figure 1-1**. **Comparing individual level changes in cortical thickness with age against the corresponding regression model output.** Each panel, referring to one brain region, shows the smoothed distribution (removing outliers for the purpose of display) of individual’s changes in cortical thickness with age-in-months [i.e., (Thickness_Y2_-Thickness_BL_) / (Age_Y2_-Age_BL_)]. The left- and right-facing arrows refer to the means of the non-outliered distributions of the left (LH) and right hemispheres (RH), respectively. The dotted line refers to an ordinate value of zero. The segmented lines refer to the unstandardized regression coefficient for age from the age-only, intracranial-volume-corrected (ICV), whole-brain-volume-corrected (WBV), and mean-cortical-thickness-corrected (SM) models.

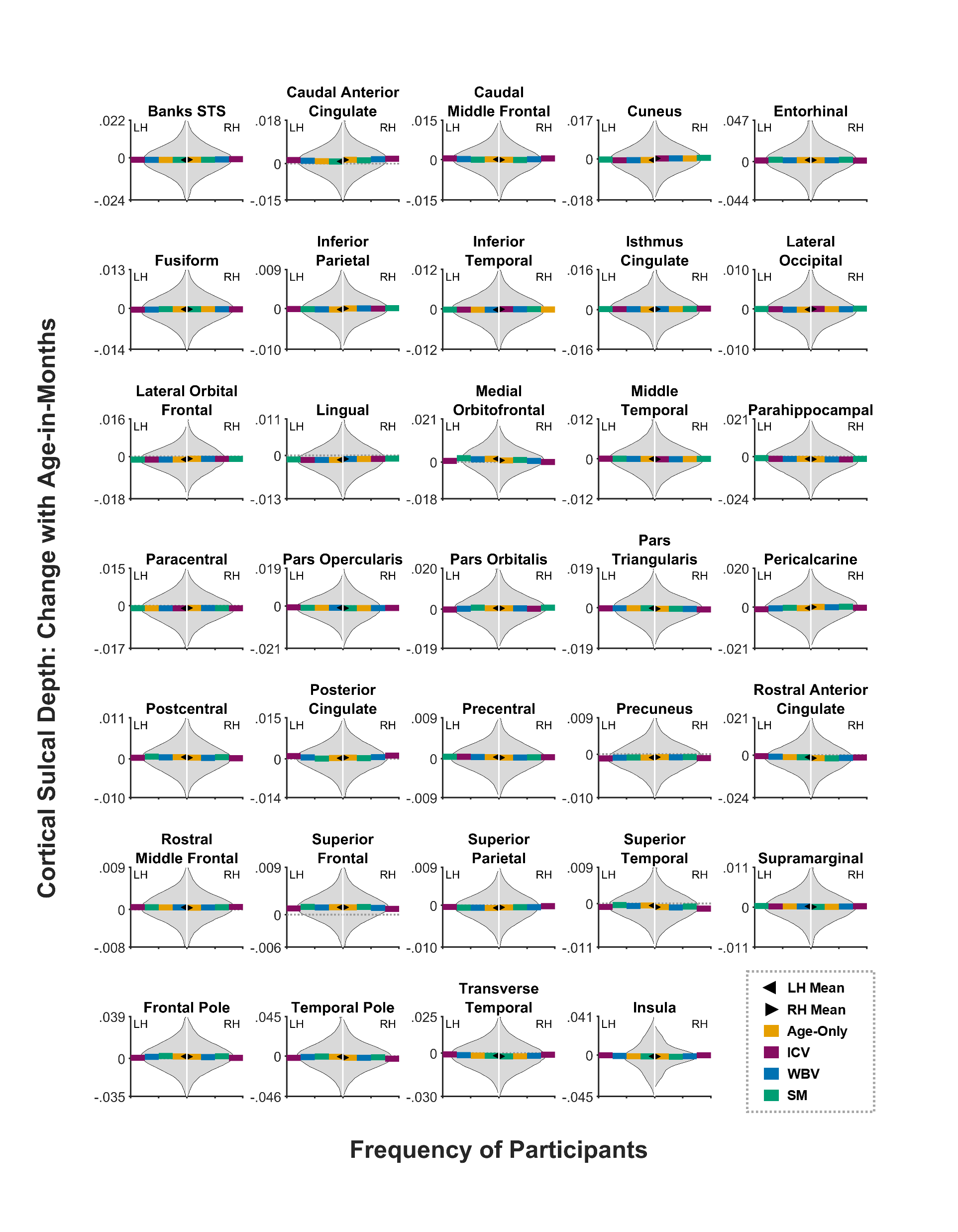

**Extended Data Figure 1-2**. **Comparing individual level changes in cortical sulcal depth with age against the corresponding regression model output.** Each panel, referring to one brain region, shows the smoothed distribution (removing outliers for the purpose of display) of individual’s changes in cortical sulcal depth with age-in-months [i.e., (Sulcal Depth_Y2_-Sulcal Depth_BL_) / (Age_Y2_-Age_BL_)]. The left- and right-facing arrows refer to the means of the non-outliered distributions of the left (LH) and right hemispheres (RH), respectively. The dotted line refers to an ordinate value of zero. The segmented lines refer to the unstandardized regression coefficient for age from the age-only, intracranial-volume-corrected (ICV), whole-brain-volume-corrected (WBV), and mean-cortical-sulcal-depth-corrected (SM) models.

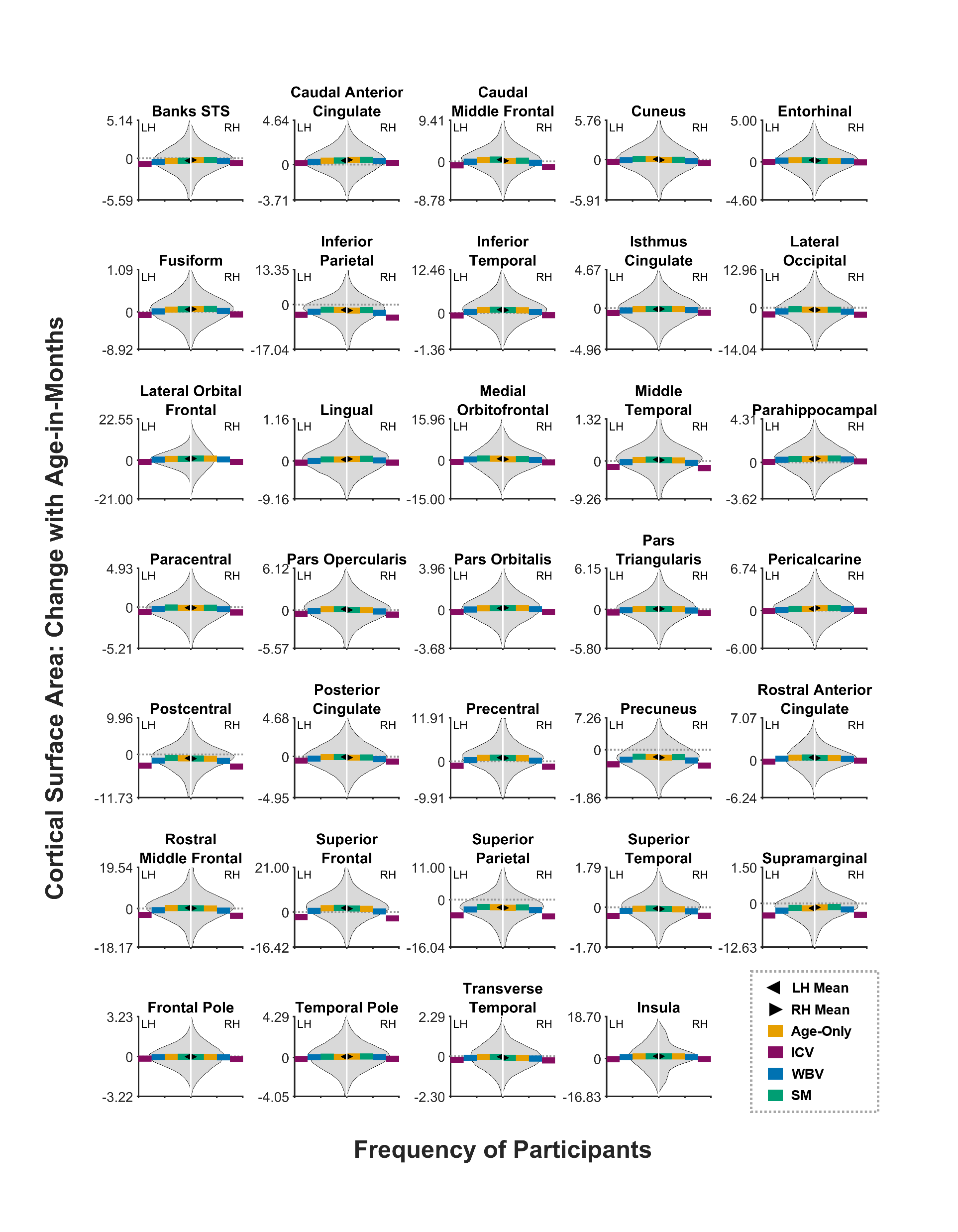

**Extended Data Figure 1-3**. **Comparing individual level changes in cortical surface area with age against the corresponding regression model output.** Each panel, referring to one brain region, shows the smoothed distribution (removing outliers for the purpose of display) of individual’s changes in cortical surface area with age-in-months [i.e., (Surface Area_Y2_-Surface Area_BL_) / (Age_Y2_-Age_BL_)]. The left- and right-facing arrows refer to the means of the non-outliered distributions of the left (LH) and right hemispheres (RH), respectively. The dotted line refers to an ordinate value of zero. The segmented lines refer to the unstandardized regression coefficient for age from the age-only, intracranial-volume-corrected (ICV), whole-brain-volume-corrected (WBV), and total-cortical-surface-area-corrected (SM) models.

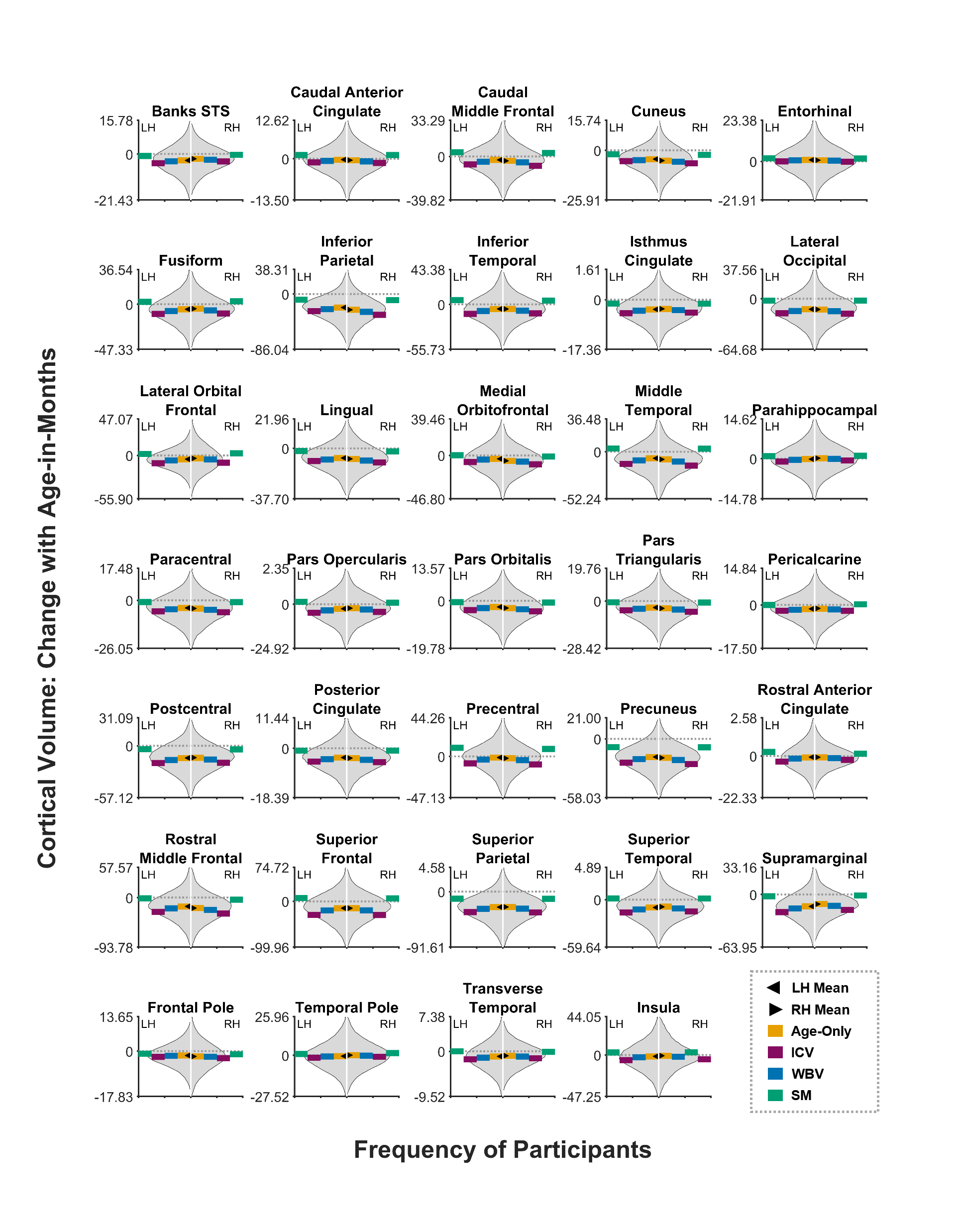

**Extended Data Figure 1-4**. **Comparing individual level changes in cortical volume with age against the corresponding regression model output.** Each panel, referring to one brain region, shows the smoothed distribution (removing outliers for the purpose of display) of individual’s changes in cortical volume with age-in-months [i.e., (Volume_Y2_-Volume_BL_) / (Age_Y2_-Age_BL_)]. The left- and right-facing arrows refer to the means of the non-outliered distributions of the left (LH) and right hemispheres (RH), respectively. The dotted line refers to an ordinate value of zero. The segmented lines refer to the unstandardized regression coefficient for age from the age-only, intracranial-volume-corrected (ICV), whole-brain-volume-corrected (WBV), and total-cortical-volume-corrected (SM) models.

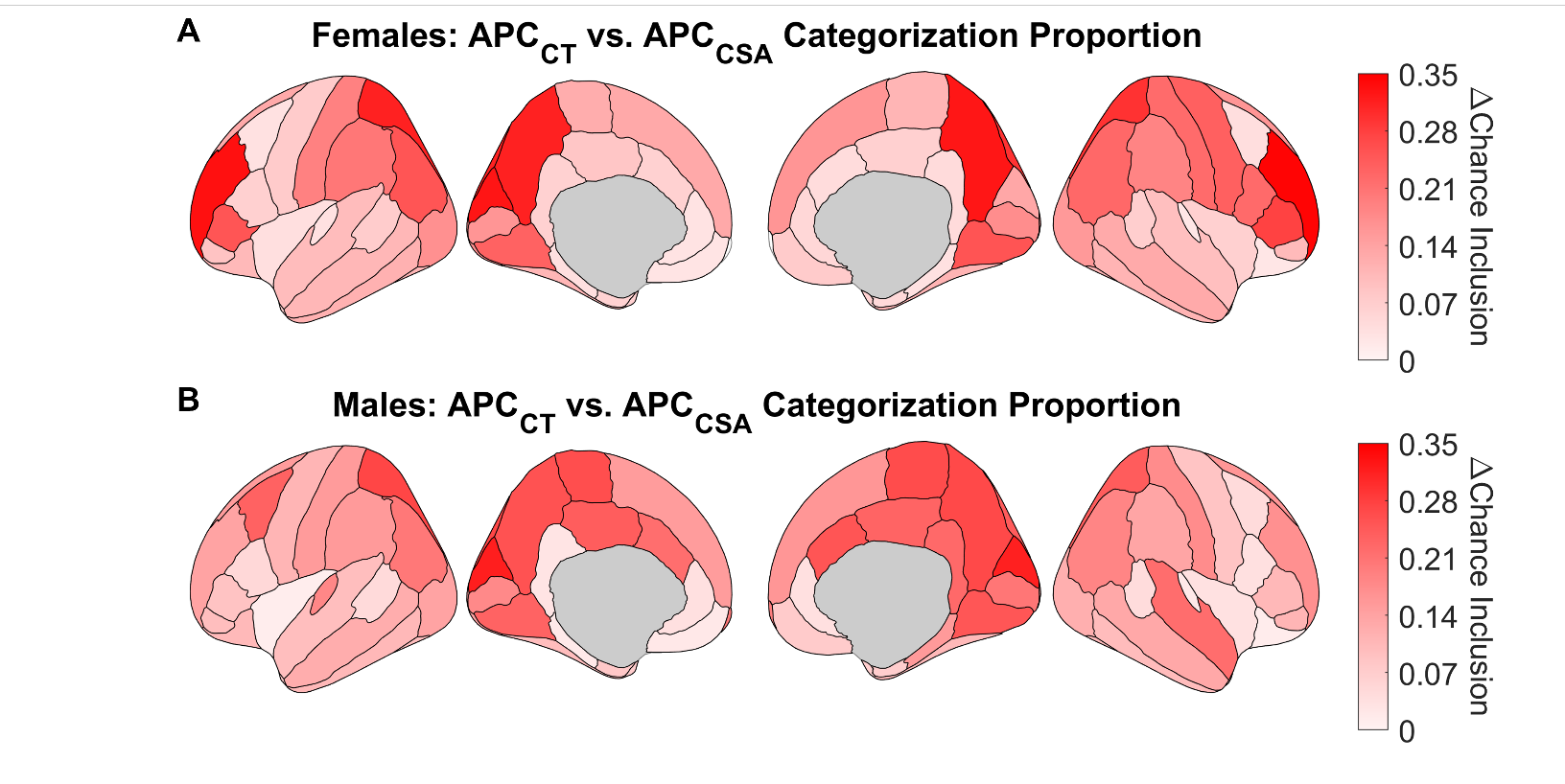

**Extended Data Figure 5-1. Proportion of individuals (relative to chance) who showed the primary group-level patterns of annual percentage change (APC) of cortical thickness (CT) and surface area (CSA) by brain region**. Each region is shaded in correspondence to how many more individuals (i.e., proportion of individuals) show the characterizing patterns of APC_CT_ vs. APC_CSA_ change by sex and brain region (see Figure 5). For example, in participants assigned female sex at birth (**A**), the left and right precuneus was primarily characterized as showing rates of cortical thinning that were faster than rates of cortical contraction (Figure 5). Accordingly, there was a 12.5% chance likelihood that any one individual would show that same patterning. Here, 43.8% and 44.9% of individuals (left and right precuneus, respectively) showed similar patterning relative to the group, which reflected a 31.3% and 32.4% increase over chance. Regions were not shaded if their corresponding proportion was not different from chance.

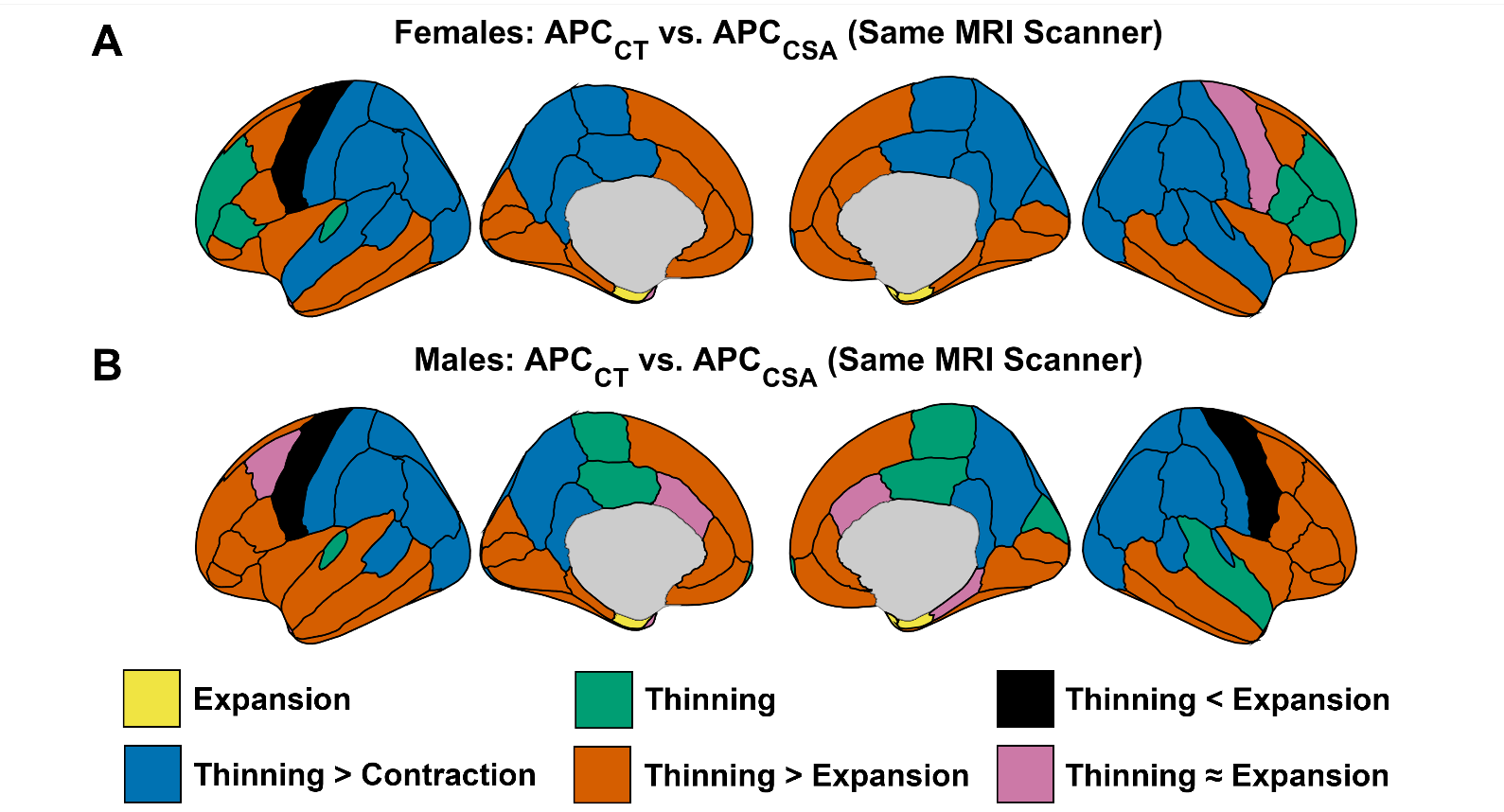

**Extended Data Figure 5-2. Regional differences in rates of developmental change in cortical thickness and surface area per annual percentage change for individuals scanned by the same MRI machine at both timepoints**. Regions are color coded with respect to differential rates of cortical thinning/thickening vs. cortical contraction/expansion, based on whether the annual percentage change rates were significantly different from 0 and whether the rate of thinning/thickening was significantly different than that of contraction/expansion. Regions that were color coded for only one type of change (i.e., “Expansion”, “Thinning”) indicates that the opponent process’s annual percentage change was not significantly different from 0.

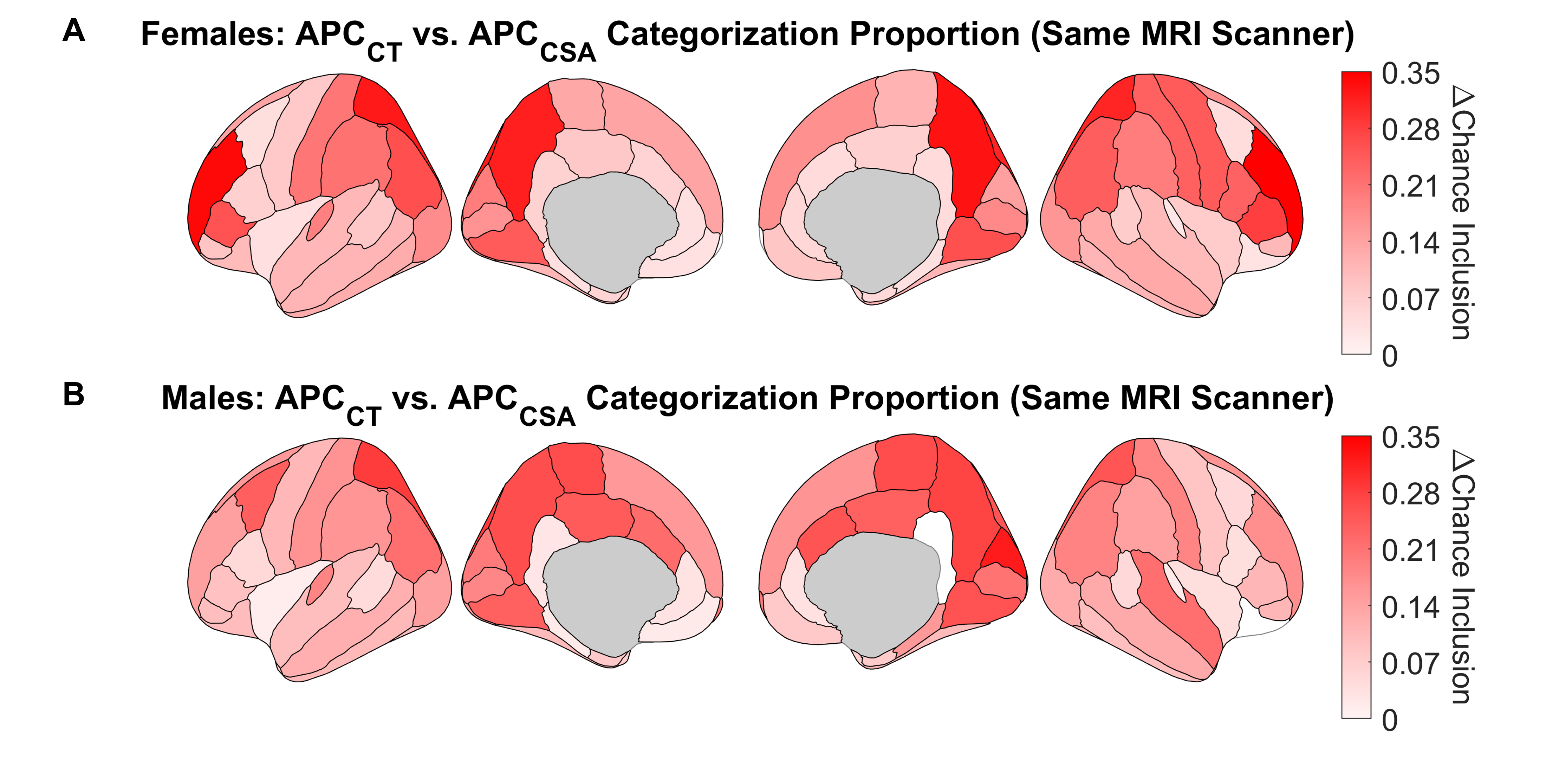

**Extended Data Figure 5-3. Proportion of individuals, who were scanned by the same MRI machine at both timepoints, (relative to chance) who showed the primary group-level patterns of annual percentage change (APC) of cortical thickness (CT) and surface area (CSA) by brain region**. Each region is shaded in correspondence to how many more individuals (i.e., proportion of individuals) show the characterizing patterns of APC_CT_ vs. APC_CSA_ change by sex and brain region. For example, in participants assigned female sex at birth (**A**), the left and right precuneus was primarily characterized as showing rates of cortical thinning that were faster than rates of cortical contraction (Extended Data Figure 5-2). Accordingly, there was a 12.5% chance likelihood that any one individual would show that same patterning. Here, 44.0% and 45.4% of individuals (left and right precuneus, respectively) showed similar patterning relative to the group, which reflected a 31.5% and 32.9% increase over chance. Regions were not shaded if their corresponding proportion was not different from chance. Note: The primary difference between this figure and Extended Data Figure 5-1 is that, for individuals assigned male sex at birth, the proportions for the right isthmus cingulate and right lateral orbital frontal cortices did not differ from chance.

**Extended Data Figure 2-1.** Associations between age (in months) and cortical thickness (mm).

| **Cortical Region (Hemisphere)** | ***n*** | ***b*** | ***t*** | ***p*** | ***r_p_*** | **95% CI (*r_p_*)** |
| --- | --- | --- | --- | --- | --- | --- |
| Banks of the Superior Temporal Sulcus (L) | 6,282 | -0.0019 | -49.78 | < .001 | -0.41 | [-.41, -.40] |
| Banks of the Superior Temporal Sulcus (R) | 6,308 | -0.0016 | -45.88 | < .001 | -0.38 | [-.39, -.37] |
| Cuneus (L) | 6,283 | -0.0022 | -63.16 | < .001 | -0.49 | [-.50, -.48] |
| Cuneus (R) | 6,287 | -0.0022 | -63.02 | < .001 | -0.49 | [-.50, -.48] |
| Fusiform (L) | 6,251 | -0.0014 | -42.89 | < .001 | -0.36 | [-.37, -.35] |
| Fusiform (R) | 6,259 | -0.0014 | -41.77 | < .001 | -0.35 | [-.36, -.34] |
| Inferior Parietal (L) | 6,220 | -0.0020 | -55.80 | < .001 | -0.45 | [-.45, -.44] |
| Inferior Parietal (R) | 6,208 | -0.0018 | -53.20 | < .001 | -0.43 | [-.44, -.42] |
| Isthmus Cingulate (L) | 6,244 | -0.0015 | -45.70 | < .001 | -0.38 | [-.39, -.37] |
| Isthmus Cingulate (R) | 6,258 | -0.0015 | -44.97 | < .001 | -0.37 | [-.38, -.37] |
| Lateral Occipital (L) | 6,296 | -0.0015 | -47.42 | < .001 | -0.39 | [-.40, -.38] |
| Lateral Occipital (R) | 6,291 | -0.0015 | -46.68 | < .001 | -0.38 | [-.39, -.38] |
| Lateral Orbital Frontal (L) | 6,258 | -0.0019 | -43.24 | < .001 | -0.36 | [-.37, -.35] |
| Lingual (L) | 6,282 | -0.0021 | -64.83 | < .001 | -0.50 | [-.51, -.49] |
| Lingual (R) | 6,284 | -0.0022 | -68.28 | < .001 | -0.52 | [-.53, -.51] |
| Medial Orbital Frontal (R) | 6,262 | -0.0021 | -43.80 | < .001 | -0.36 | [-.37, -.36] |
| Paracentral (L) | 6,254 | -0.0018 | -46.34 | < .001 | -0.38 | [-.39, -.38] |
| Paracentral (R) | 6,260 | -0.0018 | -45.46 | < .001 | -0.38 | [-.38, -.37] |
| Pars Orbitalis (L) | 6,244 | -0.0021 | -41.65 | < .001 | -0.35 | [-.36, -.34] |
| Pars Orbitalis (R) | 6,265 | -0.0021 | -44.85 | < .001 | -0.37 | [-.38, -.36] |
| Pars Triangularis (L) | 6,233 | -0.0016 | -39.69 | < .001 | -0.34 | [-.34, -.33] |
| Pars Triangularis (R) | 6,241 | -0.0016 | -40.37 | < .001 | -0.34 | [-.35, -.33] |
| Postcentral (L) | 6,237 | -0.0016 | -48.66 | < .001 | -0.40 | [-.41, -.39] |
| Postcentral (R) | 6,151 | -0.0017 | -45.51 | < .001 | -0.38 | [-.39, -.37] |
| Posterior Cingulate (L) | 6,250 | -0.0015 | -48.37 | < .001 | -0.40 | [-.40, -.39] |
| Posterior Cingulate (R) | 6,239 | -0.0015 | -48.06 | < .001 | -0.40 | [-.40, -.39] |
| Precuneus (L) | 6,256 | -0.0021 | -75.35 | < .001 | -0.56 | [-.56, -.55] |
| Precuneus (R) | 6,267 | -0.0021 | -74.04 | < .001 | -0.55 | [-.56, -.55] |
| Rostral Middle Frontal (L) | 6,225 | -0.0016 | -44.07 | < .001 | -0.37 | [-.38, -.36] |
| Rostral Middle Frontal (R) | 6,239 | -0.0019 | -48.52 | < .001 | -0.40 | [-.41, -.39] |
| Superior Parietal (L) | 6,256 | -0.0018 | -48.72 | < .001 | -0.40 | [-.41, -.39] |
| Superior Parietal (R) | 6,236 | -0.0018 | -49.25 | < .001 | -0.40 | [-.41, -.40] |
| Pericalcarine (L) | 6,302 | -0.0016 | -38.84 | < .001 | -0.33 | [-.33, -.32] |
| Supramarginal (R) | 6,148 | -0.0015 | -38.69 | < .001 | -0.33 | [-.34, -.32] |
| Superior Frontal (R) | 6,259 | -0.0015 | -38.11 | < .001 | -0.32 | [-.33, -.31] |
| Supramarginal (L) | 6,130 | -0.0015 | -37.67 | < .001 | -0.32 | [-.33, -.31] |
| Inferior Temporal (L) | 6,230 | -0.0014 | -37.38 | < .001 | -0.32 | [-.33, -.31] |
| Superior Frontal (L) | 6,245 | -0.0013 | -37.22 | < .001 | -0.32 | [-.32, -.31] |
| Lateral Orbital Frontal (R) | 6,264 | -0.0018 | -36.91 | < .001 | -0.31 | [-.32, -.31] |
| Pericalcarine (R) | 6,295 | -0.0016 | -36.11 | < .001 | -0.31 | [-.31, -.30] |
| Medial Orbital Frontal (L) | 6,259 | -0.0018 | -35.29 | < .001 | -0.30 | [-.31, -.29] |
| Inferior Temporal (R) | 6,224 | -0.0013 | -35.10 | < .001 | -0.30 | [-.31, -.29] |
| Middle Temporal (R) | 6,149 | -0.0013 | -34.22 | < .001 | -0.29 | [-.30, -.29] |
| Middle Temporal (L) | 6,169 | -0.0014 | -33.92 | < .001 | -0.29 | [-.30, -.28] |
| Caudal Anterior Cingulate (R) | 6,226 | -0.0014 | -33.74 | < .001 | -0.29 | [-.30, -.28] |
| Frontal Pole (L) | 6,225 | -0.0024 | -30.61 | < .001 | -0.26 | [-.27, -.26] |
| Superior Temporal (R) | 6,272 | -0.0011 | -30.05 | < .001 | -0.26 | [-.27, -.25] |
| Superior Temporal (L) | 6,265 | -0.0011 | -30.03 | < .001 | -0.26 | [-.27, -.25] |
| Transverse Temporal (L) | 6,288 | -0.0015 | -29.98 | < .001 | -0.26 | [-.27, -.25] |
| Pars Opercularis (R) | 6,255 | -0.0011 | -29.59 | < .001 | -0.26 | [-.26, -.25] |
| Frontal Pole (R) | 6,239 | -0.0023 | -29.43 | < .001 | -0.25 | [-.26, -.25] |
| Transverse Temporal (R) | 6,276 | -0.0016 | -28.94 | < .001 | -0.25 | [-.26, -.24] |
| Caudal Anterior Cingulate (L) | 6,277 | -0.0014 | -28.72 | < .001 | -0.25 | [-.26, -.24] |
| Rostral Anterior Cingulate (R) | 6,276 | -0.0018 | -28.42 | < .001 | -0.25 | [-.25, -.24] |
| Pars Opercularis (L) | 6,233 | -0.0010 | -28.30 | < .001 | -0.25 | [-.25, -.24] |
| Insula (L) | 6,273 | -0.0016 | -27.29 | < .001 | -0.24 | [-.25, -.23] |
| Parahippocampal (L) | 6,295 | -0.0014 | -26.71 | < .001 | -0.23 | [-.24, -.22] |
| Parahippocampal (R) | 6,287 | -0.0013 | -25.85 | < .001 | -0.22 | [-.23, -.22] |
| Insula (R) | 6,271 | -0.0015 | -24.89 | < .001 | -0.22 | [-.23, -.21] |
| Rostral Anterior Cingulate (L) | 6,283 | -0.0016 | -23.26 | < .001 | -0.20 | [-.21, -.19] |
| Caudal Middle Frontal (R) | 6,209 | -0.0010 | -22.56 | < .001 | -0.20 | [-.21, -.19] |
| Caudal Middle Frontal (L) | 6,217 | -0.0008 | -19.39 | < .001 | -0.17 | [-.18, -.16] |
| Precentral (R) | 6,022 | -0.0003 | -9.07 | < .001 | -0.08 | [-.09, -.07] |
| Precentral (L) | 6,128 | -0.0003 | -7.05 | < .001 | -0.06 | [-.07, -.05] |
| Temporal Pole (L) | 6,192 | -0.0006 | -5.45 | < .001 | -0.05 | [-.06, -.04] |
| Entorhinal (R) | 6,248 | 0.0003 | 2.26 | .024 | 0.02 | [.01, .03] |
| Entorhinal (L) | 6,227 | 0.0002 | 1.85 | .065 | 0.02 | [.01, .03] |
| Temporal Pole (R) | 6,156 | -0.0001 | -0.81 | .416 | -0.01 | [-.02, .002] |

*Note*. Cortical regions are sorted vertically by *p*-value (uncorrected) for associations with age. Analyses included fixed effects of sex, age, age^2^, Sex × Age, Sex × Age^2^, and whole-brain volume. The random-effects structure included random intercepts of participant ID and magnetic resonance imaging (MRI) scanner serial number. Associations that passed false-discovery-rate (FDR) correction are shaded. *n* = number of participants included in analysis of that region. *t* = *t*-statistic. *b* = unstandardized regression coefficient [i.e., change in cortical thickness (mm) with a one-month increase in age]. 95% CI = 95% confidence interval. *r_p_* = partial correlation coefficient (i.e., effect size).

**Extended Data Figure 2-2.** Associations between age (in months) and sulcal depth (mm).

| **Cortical Region (Hemisphere)** | ***n*** | ***b*** | ***t*** | ***p*** | ***r_p_*** | **95% CI (*r_p_*)** |
| --- | --- | --- | --- | --- | --- | --- |
| Superior Frontal (L) | 6,294 | 0.0013 | 35.80 | < .001 | 0.30 | [.30, .31] |
| Superior Frontal (R) | 6,293 | 0.0012 | 30.95 | < .001 | 0.27 | [.26, .27] |
| Lingual (L) | 6,271 | -0.0014 | -24.12 | < .001 | -0.21 | [-.22, -.20] |
| Caudal Anterior Cingulate (R) | 6,262 | 0.0018 | 23.03 | < .001 | 0.20 | [.19, .21] |
| Superior Temporal (R) | 6,278 | -0.0010 | -21.82 | < .001 | -0.19 | [-.20, -.18] |
| Lateral Orbital Frontal (L) | 6,284 | -0.0013 | -20.54 | < .001 | -0.18 | [-.19, -.17] |
| Lingual (R) | 6,272 | -0.0011 | -18.67 | < .001 | -0.16 | [-.17, -.16] |
| Precuneus (L) | 6,299 | -0.0009 | -17.93 | < .001 | -0.16 | [-.17, -.15] |
| Caudal Anterior Cingulate (L) | 6,270 | 0.0012 | 15.50 | < .001 | 0.14 | [.13, .15] |
| Parahippocampal (R) | 6,294 | -0.0016 | -14.99 | < .001 | -0.13 | [-.14, -.12] |
| Medial Orbital Frontal (L) | 6,294 | 0.0013 | 14.92 | < .001 | 0.13 | [.12, .14] |
| Transverse Temporal (R) | 6,295 | -0.0018 | -14.70 | < .001 | -0.13 | [-.14, -.12] |
| Precuneus (R) | 6,291 | -0.0008 | -14.25 | < .001 | -0.13 | [-.13, -.12] |
| Rostral Anterior Cingulate (R) | 6,282 | -0.0016 | -14.14 | < .001 | -0.13 | [-.13, -.12] |
| Superior Temporal (L) | 6,266 | -0.0006 | -13.41 | < .001 | -0.12 | [-.13, -.11] |
| Paracentral (L) | 6,257 | -0.0010 | -13.04 | < .001 | -0.12 | [-.12, -.11] |
| Lateral Orbital Frontal (R) | 6,283 | -0.0010 | -12.51 | < .001 | -0.11 | [-.12, -.10] |
| Parahippocampal (L) | 6,310 | -0.0013 | -12.08 | < .001 | -0.11 | [-.12, -.10] |
| Superior Parietal (L) | 6,289 | -0.0006 | -12.06 | < .001 | -0.11 | [-.12, -.10] |
| Transverse Temporal (L) | 6,306 | -0.0014 | -11.82 | < .001 | -0.10 | [-.11, -.10] |
| Paracentral (R) | 6,267 | -0.0009 | -11.36 | < .001 | -0.10 | [-.11, -.09] |
| Banks of the Superior Temporal Sulcus (L) | 6,150 | -0.0012 | -10.44 | < .001 | -0.09 | [-.10, -.08] |
| Pars Opercularis (L) | 6,225 | -0.0009 | -10.34 | < .001 | -0.09 | [-.10, -.08] |
| Rostral Middle Frontal (L) | 6,282 | 0.0005 | 9.94 | < .001 | 0.09 | [.08, .10] |
| Rostral Anterior Cingulate (L) | 6,298 | -0.0010 | -9.63 | < .001 | -0.09 | [-.09, -.08] |
| Temporal Pole (R) | 6,264 | -0.0019 | -9.11 | < .001 | -0.08 | [-.09, -.07] |
| Banks of the Superior Temporal Sulcus (R) | 6,111 | -0.0010 | -9.03 | < .001 | -0.08 | [-.09, -.07] |
| Posterior Cingulate (R) | 6,287 | 0.0007 | 8.97 | < .001 | 0.08 | [.07, .09] |
| Precentral (L) | 6,218 | 0.0004 | 8.57 | < .001 | 0.08 | [.07, .09] |
| Insula (R) | 6,278 | -0.0013 | -8.04 | < .001 | -0.07 | [-.08, -.06] |
| Pars Opercularis (R) | 6,169 | -0.0010 | -7.87 | < .001 | -0.07 | [-.08, -.06] |
| Posterior Cingulate (L) | 6,278 | 0.0005 | 7.87 | < .001 | 0.07 | [.06, .08] |
| Frontal Pole (L) | 6,238 | 0.0014 | 7.74 | < .001 | 0.07 | [.06, .08] |
| Medial Orbital Frontal (R) | 6,278 | 0.0006 | 7.69 | < .001 | 0.07 | [.06, .08] |
| Postcentral (L) | 6,252 | 0.0004 | 7.66 | < .001 | 0.07 | [.06, .08] |
| Rostral Middle Frontal (R) | 6,292 | 0.0004 | 7.66 | < .001 | 0.07 | [.06, .08] |
| Superior Parietal (R) | 6,269 | -0.0005 | -7.59 | < .001 | -0.07 | [-.08, -.06] |
| Pars Triangularis (R) | 6,260 | -0.0006 | -7.27 | < .001 | -0.06 | [-.07, -.06] |
| Entorhinal (L) | 6,312 | 0.0016 | 7.01 | < .001 | 0.06 | [.05, .07] |
| Entorhinal (R) | 6,310 | 0.0016 | 6.86 | < .001 | 0.06 | [.05, .07] |
| Cuneus (L) | 6,290 | -0.0006 | -6.69 | < .001 | -0.06 | [-.07, -.05] |
| Pericalcarine (L) | 6,219 | -0.0006 | -6.68 | < .001 | -0.06 | [-.07, -.05] |
| Frontal Pole (R) | 6,264 | 0.0012 | 6.65 | < .001 | 0.06 | [.05, .07] |
| Fusiform (L) | 6,292 | -0.0004 | -5.94 | < .001 | -0.05 | [-.06, -.04] |
| Insula (L) | 6,311 | -0.0011 | -5.75 | < .001 | -0.05 | [-.06, -.04] |
| Precentral (R) | 6,235 | 0.0002 | 4.81 | < .001 | 0.04 | [.03, .05] |
| Postcentral (R) | 6,178 | 0.0003 | 4.78 | < .001 | 0.04 | [.03, .05] |
| Fusiform (R) | 6,292 | -0.0003 | -4.72 | < .001 | -0.04 | [-.05, -.03] |
| Temporal Pole (L) | 6,260 | -0.0011 | -4.58 | < .001 | -0.04 | [-.05, -.03] |
| Supramarginal (L) | 6,264 | 0.0002 | 3.91 | < .001 | 0.03 | [.03, .04] |
| Pars Orbitalis (R) | 6,284 | 0.0004 | 3.84 | < .001 | 0.03 | [.03, .04] |
| Inferior Parietal (L) | 6,283 | -0.0002 | -3.79 | < .001 | -0.03 | [-.04, -.02] |
| Inferior Temporal (L) | 6,290 | -0.0002 | -3.70 | < .001 | -0.03 | [-.04, -.02] |
| Lateral Occipital (L) | 6,304 | -0.0002 | -3.56 | < .001 | -0.03 | [-.04, -.02] |
| Caudal Middle Frontal (L) | 6,228 | 0.0002 | 3.00 | .003 | 0.03 | [.02, .04] |
| Isthmus Cingulate (R) | 6,254 | 0.0002 | 2.82 | .005 | 0.03 | [.02, .03] |
| Pars Triangularis (L) | 6,253 | -0.0002 | -2.62 | .009 | -0.02 | [-.03, -.01] |
| Pars Orbitalis (L) | 6,285 | 0.0002 | 2.52 | .012 | 0.02 | [.01, .03] |
| Middle Temporal (R) | 6,287 | -0.0001 | -2.45 | .014 | -0.02 | [-.03, -.01] |
| Supramarginal (R) | 6,269 | 0.0002 | 2.43 | .015 | 0.02 | [.01, .03] |
| Cuneus (R) | 6,296 | 0.0002 | 2.30 | .022 | 0.02 | [.01, .03] |
| Caudal Middle Frontal (R) | 6,217 | 0.0002 | 1.88 | .061 | 0.02 | [.01, .03] |
| Isthmus Cingulate (L) | 6,252 | 0.0001 | 1.78 | .074 | 0.02 | [.01, .02] |
| Inferior Temporal (R) | 6,285 | -0.0001 | -1.62 | .105 | -0.01 | [-.02, -.01] |
| Lateral Occipital (R) | 6,302 | -0.0001 | -1.52 | .129 | -0.01 | [-.02, -.005] |
| Inferior Parietal (R) | 6,291 | 0.0000 | 0.60 | .546 | 0.01 | [-.004, .01] |
| Middle Temporal (L) | 6,276 | 0.0000 | -0.51 | .608 | 0.00 | [-.01, .004] |
| Pericalcarine (R) | 6,227 | 0.0000 | 0.17 | .869 | 0.00 | [-.01, .01] |

*Note*. Cortical regions are sorted vertically by *p*-value (uncorrected) for associations with age. Analyses included fixed effects of sex, age, age^2^, Sex × Age, Sex × Age^2^, and whole-brain volume. The random-effects structure included random intercepts of participant ID and magnetic resonance imaging (MRI) scanner serial number. Associations that passed false-discovery-rate (FDR) correction are shaded. *n* = number of participants included in analysis of that region. *t* = *t*-statistic. *b* = unstandardized regression coefficient [i.e., change in sulcal depth (mm) with a one-month increase in age]. 95% CI = 95% confidence interval. *r_p_* = partial correlation coefficient (i.e., effect size).

**Extended Data Figure 2-3.** Associations between age (in months) and cortical surface area (mm^2^).

| **Cortical Region (Hemisphere)** | ***n*** | ***b*** | ***t*** | ***p*** | ***r_p_*** | **95% CI (*r_p_*)** |
| --- | --- | --- | --- | --- | --- | --- |
| Precuneus (L) | 6,282 | -1.5764 | -25.03 | < .001 | -0.22 | [-.23, -.21] |
| Inferior Parietal (L) | 6,296 | -1.9705 | -24.87 | < .001 | -0.22 | [-.22, -.21] |
| Precuneus (R) | 6,271 | -1.6953 | -24.65 | < .001 | -0.22 | [-.22, -.21] |
| Caudal Anterior Cingulate (L) | 6,201 | 0.4177 | 22.57 | < .001 | 0.20 | [.19, .21] |
| Caudal Anterior Cingulate (R) | 6,256 | 0.5016 | 22.43 | < .001 | 0.20 | [.19, .21] |
| Inferior Parietal (R) | 6,299 | -2.1522 | -21.28 | < .001 | -0.19 | [-.19, -.18] |
| Parahippocampal (R) | 6,222 | 0.3739 | 20.69 | < .001 | 0.18 | [.17, .19] |
| Superior Parietal (L) | 6,261 | -2.4410 | -19.77 | < .001 | -0.17 | [-.18, -.17] |
| Superior Parietal (R) | 6,267 | -2.6024 | -18.86 | < .001 | -0.17 | [-.17, -.16] |
| Parahippocampal (L) | 6,196 | 0.3284 | 17.50 | < .001 | 0.16 | [.15, .16] |
| Inferior Temporal (L) | 6,317 | 0.9412 | 16.51 | < .001 | 0.15 | [.14, .15] |
| Inferior Temporal (R) | 6,313 | 0.8567 | 16.40 | < .001 | 0.14 | [.14, .15] |
| Fusiform (R) | 6,305 | 0.6819 | 16.18 | < .001 | 0.14 | [.13, .15] |
| Superior Frontal (L) | 6,286 | 1.9567 | 15.95 | < .001 | 0.14 | [.13, .15] |
| Rostral Anterior Cingulate (R) | 6,313 | 0.3445 | 15.27 | < .001 | 0.13 | [.13, .14] |
| Postcentral (L) | 6,239 | -0.9923 | -14.18 | < .001 | -0.13 | [-.13, -.12] |
| Rostral Anterior Cingulate (L) | 6,301 | 0.4440 | 14.14 | < .001 | 0.13 | [.12, .13] |
| Postcentral (R) | 6,237 | -1.1041 | -13.82 | < .001 | -0.12 | [-.13, -.11] |
| Fusiform (L) | 6,298 | 0.5950 | 13.73 | < .001 | 0.12 | [.11, .13] |
| Pericalcarine (R) | 6,309 | 0.4322 | 13.22 | < .001 | 0.12 | [.11, .13] |
| Lateral Orbital Frontal (L) | 6,305 | 0.7776 | 12.94 | < .001 | 0.11 | [.11, .12] |
| Precentral (L) | 6,250 | 1.0182 | 12.41 | < .001 | 0.11 | [.10, .12] |
| Insula (L) | 6,266 | 0.9519 | 12.30 | < .001 | 0.11 | [.10, .12] |
| Supramarginal (L) | 6,276 | -1.2877 | -12.12 | < .001 | -0.11 | [-.12, -.10] |
| Insula (R) | 6,267 | 0.9382 | 11.56 | < .001 | 0.10 | [.09, .11] |
| Lingual (R) | 6,296 | 0.5180 | 11.11 | < .001 | 0.10 | [.09, .11] |
| Banks of the Superior Temporal Sulcus (L) | 6,245 | -0.3051 | -10.28 | < .001 | -0.09 | [-.10, -.08] |
| Superior Frontal (R) | 6,277 | 1.6240 | 10.17 | < .001 | 0.09 | [.08, .10] |
| Pars Orbitalis (R) | 6,307 | 0.1844 | 9.94 | < .001 | 0.09 | [.08, .10] |
| Lateral Orbital Frontal (R) | 6,291 | 0.9190 | 9.88 | < .001 | 0.09 | [.08, .10] |
| Transverse Temporal (R) | 6,232 | -0.0838 | -9.84 | < .001 | -0.09 | [-.10, -.08] |
| Supramarginal (R) | 6,266 | -1.0497 | -9.36 | < .001 | -0.08 | [-.09, -.07] |
| Pericalcarine (L) | 6,307 | 0.2810 | 9.22 | < .001 | 0.08 | [.07, .09] |
| Lateral Occipital (R) | 6,308 | -0.6377 | -9.19 | < .001 | -0.08 | [-.09, -.07] |
| Precentral (R) | 6,243 | 0.8876 | 9.15 | < .001 | 0.08 | [.07, .09] |
| Banks of the Superior Temporal Sulcus (R) | 6,267 | -0.2037 | -8.85 | < .001 | -0.08 | [-.09, -.07] |
| Pars Orbitalis (L) | 6,312 | 0.1391 | 8.73 | < .001 | 0.08 | [.07, .09] |
| Superior Temporal (R) | 6,255 | -0.4541 | -8.66 | < .001 | -0.08 | [-.09, -.07] |
| Medial Orbital Frontal (L) | 6,276 | 0.5866 | 8.57 | < .001 | 0.08 | [.07, .09] |
| Lingual (L) | 6,289 | 0.3661 | 8.36 | < .001 | 0.07 | [.07, .08] |
| Lateral Occipital (L) | 6,313 | -0.5160 | -7.95 | < .001 | -0.07 | [-.08, -.06] |
| Entorhinal (L) | 6,246 | 0.1856 | 7.73 | < .001 | 0.07 | [.06, .08] |
| Caudal Middle Frontal (L) | 6,283 | 0.4302 | 6.93 | < .001 | 0.06 | [.05, .07] |
| Medial Orbital Frontal (R) | 6,290 | 0.3802 | 6.67 | < .001 | 0.06 | [.05, .07] |
| Temporal Pole (R) | 6,243 | 0.1161 | 6.43 | < .001 | 0.06 | [.05, .07] |
| Entorhinal (R) | 6,225 | 0.1170 | 5.60 | < .001 | 0.05 | [.04, .06] |
| Middle Temporal (L) | 6,291 | 0.3306 | 5.55 | < .001 | 0.05 | [.04, .06] |
| Temporal Pole (L) | 6,274 | 0.0997 | 5.26 | < .001 | 0.05 | [.04, .06] |
| Superior Temporal (L) | 6,266 | -0.3168 | -5.16 | < .001 | -0.05 | [-.05, -.04] |
| Posterior Cingulate (R) | 6,265 | -0.1146 | -4.75 | < .001 | -0.04 | [-.05, -.03] |
| Isthmus Cingulate (L) | 6,222 | -0.1138 | -4.71 | < .001 | -0.04 | [-.05, -.03] |
| Isthmus Cingulate (R) | 6,235 | -0.1027 | -4.19 | < .001 | -0.04 | [-.05, -.03] |
| Pars Opercularis (L) | 6,226 | 0.1503 | 4.03 | < .001 | 0.04 | [.03, .05] |
| Middle Temporal (R) | 6,289 | 0.2176 | 3.83 | < .001 | 0.03 | [.03, .04] |
| Cuneus (L) | 6,289 | 0.0886 | 3.38 | .001 | 0.03 | [.02, .04] |
| Pars Triangularis (L) | 6,298 | 0.0799 | 2.60 | .009 | 0.02 | [.01, .03] |
| Posterior Cingulate (L) | 6,251 | -0.0592 | -2.59 | .010 | -0.02 | [-.03, -.01] |
| Transverse Temporal (L) | 6,209 | -0.0279 | -2.57 | .010 | -0.02 | [-.03, -.01] |
| Caudal Middle Frontal (R) | 6,277 | 0.1830 | 2.34 | .019 | 0.02 | [.01, .03] |
| Rostral Middle Frontal (L) | 6,308 | 0.2861 | 2.32 | .021 | 0.02 | [.01, .03] |
| Paracentral (R) | 6,242 | -0.0662 | -2.17 | .030 | -0.02 | [-.03, -.01] |
| Paracentral (L) | 6,256 | -0.0541 | -2.06 | .039 | -0.02 | [-.03, -.01] |
| Pars Triangularis (R) | 6,269 | 0.0811 | 1.86 | .063 | 0.02 | [.01, .03] |
| Frontal Pole (R) | 6,277 | -0.0234 | -1.66 | .097 | -0.01 | [-.02, -.01] |
| Frontal Pole (L) | 6,289 | -0.0154 | -1.41 | .160 | -0.01 | [-.02, -.004] |
| Pars Opercularis (R) | 6,234 | 0.0356 | 0.94 | .348 | 0.01 | [-.001, .02] |
| Rostral Middle Frontal (R) | 6,308 | 0.1362 | 0.90 | .368 | 0.01 | [-.001, .02] |
| Cuneus (R) | 6,284 | -0.0231 | -0.77 | .442 | -0.01 | [-.02, .002] |

*Note*. Cortical regions are sorted vertically by *p*-value (uncorrected) for associations with age. Analyses included fixed effects of sex, age, age^2^, Sex × Age, Sex × Age^2^, and total surface area. The random-effects structure included random intercepts of participant ID and magnetic resonance imaging (MRI) scanner serial number. Associations that passed false-discovery-rate (FDR) correction are shaded. *n* = number of participants included in analysis of that region. *t* = *t*-statistic. *b* = unstandardized regression coefficient [i.e., change in cortical surface area (mm^2^) with a one-month increase in age]. 95% CI = 95% confidence interval. *r_p_* = partial correlation coefficient (i.e., effect size).

**Extended Data Figure 2-4.** Associations between age (in months) and cortical volume (mm^3^).

| **Cortical Region (Hemisphere)** | ***n*** | ***b*** | ***t*** | ***p*** | ***r_p_*** | **95% CI (*r_p_*)** |
| --- | --- | --- | --- | --- | --- | --- |
| Cuneus (L) | 6,256 | -5.1124 | -63.89 | < .001 | -0.50 | [-.50, -.49] |
| Cuneus (R) | 6,249 | -5.9495 | -59.24 | < .001 | -0.47 | [-.48, -.46] |
| Inferior Parietal (L) | 6,297 | -23.5929 | -93.60 | < .001 | -0.64 | [-.65, -.64] |
| Inferior Parietal (R) | 6,293 | -27.7648 | -86.61 | < .001 | -0.61 | [-.62, -.61] |
| Isthmus Cingulate (L) | 6,239 | -3.8674 | -56.13 | < .001 | -0.45 | [-.46, -.44] |
| Isthmus Cingulate (R) | 6,238 | -3.6428 | -52.13 | < .001 | -0.42 | [-.43, -.42] |
| Lateral Occipital (L) | 6,300 | -15.5763 | -69.85 | < .001 | -0.53 | [-.53, -.52] |
| Lateral Occipital (R) | 6,297 | -15.9988 | -67.90 | < .001 | -0.52 | [-.52, -.51] |
| Lingual (L) | 6,288 | -8.3829 | -67.01 | < .001 | -0.51 | [-.52, -.51] |
| Lingual (R) | 6,291 | -9.1830 | -69.24 | < .001 | -0.53 | [-.53, -.52] |
| Middle Temporal (L) | 6,296 | -9.5701 | -44.01 | < .001 | -0.37 | [-.37, -.36] |
| Middle Temporal (R) | 6,303 | -10.9110 | -51.10 | < .001 | -0.41 | [-.42, -.41] |
| Paracentral (L) | 6,243 | -4.7940 | -48.98 | < .001 | -0.40 | [-.41, -.39] |
| Paracentral (R) | 6,243 | -5.1692 | -45.86 | < .001 | -0.38 | [-.39, -.37] |
| Pars Orbitalis (L) | 6,300 | -3.0290 | -44.01 | < .001 | -0.37 | [-.37, -.36] |
| Pars Orbitalis (R) | 6,302 | -3.5527 | -46.51 | < .001 | -0.38 | [-.39, -.38] |
| Pars Triangularis (L) | 6,279 | -4.6620 | -42.80 | < .001 | -0.36 | [-.36, -.35] |
| Postcentral (L) | 6,263 | -15.5740 | -74.81 | < .001 | -0.56 | [-.56, -.55] |
| Postcentral (R) | 6,276 | -15.0806 | -70.33 | < .001 | -0.53 | [-.54, -.53] |
| Posterior Cingulate (L) | 6,265 | -4.0713 | -57.53 | < .001 | -0.46 | [-.46, -.45] |
| Posterior Cingulate (R) | 6,267 | -4.2937 | -60.90 | < .001 | -0.48 | [-.48, -.47] |
| Precuneus (L) | 6,297 | -19.9179 | -97.25 | < .001 | -0.65 | [-.66, -.65] |
| Precuneus (R) | 6,272 | -20.7745 | -93.62 | < .001 | -0.64 | [-.65, -.64] |
| Rostral Middle Frontal (L) | 6,300 | -20.3272 | -49.96 | < .001 | -0.41 | [-.41, -.40] |
| Rostral Middle Frontal (R) | 6,303 | -23.2969 | -48.59 | < .001 | -0.40 | [-.40, -.39] |
| Superior Frontal (L) | 6,296 | -19.8167 | -46.89 | < .001 | -0.39 | [-.39, -.38] |
| Superior Parietal (L) | 6,270 | -28.2911 | -72.68 | < .001 | -0.54 | [-.55, -.54] |
| Superior Parietal (R) | 6,264 | -28.1238 | -63.90 | < .001 | -0.50 | [-.50, -.49] |
| Superior Temporal (L) | 6,282 | -12.2960 | -58.92 | < .001 | -0.47 | [-.47, -.46] |
| Superior Temporal (R) | 6,270 | -11.0213 | -60.55 | < .001 | -0.48 | [-.48, -.47] |
| Supramarginal (L) | 6,287 | -16.9548 | -49.38 | < .001 | -0.40 | [-.41, -.40] |
| Supramarginal (R) | 6,282 | -13.8028 | -38.06 | < .001 | -0.32 | [-.33, -.31] |
| Lateral Orbital Frontal (L) | 6,293 | -6.4008 | -37.76 | < .001 | -0.32 | [-.33, -.31] |
| Medial Orbital Frontal (R) | 6,280 | -6.6740 | -37.49 | < .001 | -0.32 | [-.33, -.31] |
| Superior Frontal (R) | 6,288 | -19.5608 | -37.30 | < .001 | -0.32 | [-.32, -.31] |
| Inferior Temporal (R) | 6,308 | -7.9110 | -37.04 | < .001 | -0.31 | [-.32, -.31] |
| Fusiform (L) | 6,288 | -7.0749 | -37.01 | < .001 | -0.31 | [-.32, -.31] |
| Fusiform (R) | 6,295 | -6.4783 | -36.07 | < .001 | -0.31 | [-.31, -.30] |
| Banks of the Superior Temporal Sulcus (L) | 6,225 | -3.4401 | -35.61 | < .001 | -0.30 | [-.31, -.30] |
| Banks of the Superior Temporal Sulcus (R) | 6,254 | -2.6564 | -35.39 | < .001 | -0.30 | [-.31, -.29] |
| Inferior Temporal (L) | 6,309 | -8.1219 | -34.86 | < .001 | -0.30 | [-.30, -.29] |
| Transverse Temporal (R) | 6,207 | -1.1035 | -34.24 | < .001 | -0.29 | [-.30, -.29] |
| Transverse Temporal (L) | 6,256 | -1.3172 | -33.91 | < .001 | -0.29 | [-.30, -.28] |
| Pars Triangularis (R) | 6,262 | -5.1084 | -33.25 | < .001 | -0.28 | [-.29, -.28] |
| Frontal Pole (L) | 6,262 | -1.8638 | -31.73 | < .001 | -0.27 | [-.28, -.26] |
| Frontal Pole (R) | 6,246 | -2.2569 | -30.73 | < .001 | -0.27 | [-.27, -.26] |
| Pericalcarine (L) | 6,296 | -1.9538 | -28.93 | < .001 | -0.25 | [-.26, -.24] |
| Pars Opercularis (L) | 6,227 | -3.5425 | -27.87 | < .001 | -0.24 | [-.25, -.23] |
| Medial Orbital Frontal (L) | 6,280 | -4.4796 | -23.37 | < .001 | -0.20 | [-.21, -.20] |
| Pericalcarine (R) | 6,287 | -1.8015 | -23.02 | < .001 | -0.20 | [-.21, -.19] |
| Caudal Middle Frontal (L) | 6,294 | -4.7220 | -22.90 | < .001 | -0.20 | [-.21, -.19] |
| Lateral Orbital Frontal (R) | 6,277 | -5.2328 | -22.75 | < .001 | -0.20 | [-.21, -.19] |
| Pars Opercularis (R) | 6,232 | -2.9934 | -22.22 | < .001 | -0.20 | [-.20, -.19] |
| Caudal Middle Frontal (R) | 6,284 | -5.3800 | -21.06 | < .001 | -0.18 | [-.19, -.18] |
| Precentral (R) | 6,229 | -4.3700 | -15.05 | < .001 | -0.13 | [-.14, -.12] |
| Precentral (L) | 6,276 | -3.6380 | -14.84 | < .001 | -0.13 | [-.14, -.12] |
| Rostral Anterior Cingulate (R) | 6,313 | -1.1556 | -13.90 | < .001 | -0.12 | [-.13, -.11] |
| Insula (L) | 6,272 | -2.6987 | -13.18 | < .001 | -0.12 | [-.13, -.11] |
| Rostral Anterior Cingulate (L) | 6,298 | -1.4033 | -12.98 | < .001 | -0.11 | [-.12, -.11] |
| Caudal Anterior Cingulate (R) | 6,241 | -0.8953 | -12.33 | < .001 | -0.11 | [-.12, -.10] |
| Caudal Anterior Cingulate (L) | 6,209 | -0.7235 | -10.83 | < .001 | -0.10 | [-.11, -.09] |
| Insula (R) | 6,267 | -1.9534 | -8.96 | < .001 | -0.08 | [-.09, -.07] |
| Temporal Pole (L) | 6,263 | -0.8502 | -6.58 | < .001 | -0.06 | [-.07, -.05] |
| Parahippocampal (L) | 6,221 | -0.4272 | -5.99 | < .001 | -0.05 | [-.06, -.04] |
| Entorhinal (L) | 6,139 | 0.5576 | 5.15 | < .001 | 0.05 | [.04, .06] |
| Entorhinal (R) | 6,163 | 0.4137 | 3.83 | < .001 | 0.03 | [.03, .04] |
| Temporal Pole (R) | 6,245 | -0.2430 | -1.97 | .049 | -0.02 | [-.03, -.01] |
| Parahippocampal (R) | 6,229 | -0.0674 | -1.00 | .319 | -0.01 | [-.02, .00002] |

*Note*. Cortical regions are sorted vertically by *p*-value (uncorrected) for associations with age. Analyses included fixed effects of sex, age, age^2^, Sex × Age, Sex × Age^2^, and whole-brain volume. The random-effects structure included random intercepts of participant ID and magnetic resonance imaging (MRI) scanner serial number. Associations that passed false-discovery-rate (FDR) correction are shaded. *n* = number of participants included in analysis of that region. *t* = *t*-statistic. *b* = unstandardized regression coefficient [i.e., change in cortical volume (mm^3^) with a one-month increase in age]. 95% CI = 95% confidence interval. *r_p_* = partial correlation coefficient (i.e., effect size).

**Extended Data Figure 3-1.** Sex × Age (in months) interactions on cortical thickness (mm).

| **Cortical Region (Hemisphere)** | ***n*** | ***b*** | ***t*** | ***p*** | ***r_p_*** | **95% CI (*r_p_*)** |
| --- | --- | --- | --- | --- | --- | --- |
| Precuneus (R) | 6,267 | -0.0002 | -8.36 | < .001 | -0.07 | [-.08, -.07] |
| Isthmus Cingulate (L) | 6,244 | -0.0003 | -8.07 | < .001 | -0.07 | [-.08, -.06] |
| Precuneus (L) | 6,256 | -0.0002 | -7.83 | < .001 | -0.07 | [-.08, -.06] |
| Isthmus Cingulate (R) | 6,258 | -0.0003 | -7.67 | < .001 | -0.07 | [-.08, -.06] |
| Inferior Parietal (R) | 6,208 | -0.0003 | -7.59 | < .001 | -0.07 | [-.08, -.06] |
| Inferior Parietal (L) | 6,220 | -0.0002 | -7.17 | < .001 | -0.06 | [-.07, -.06] |
| Posterior Cingulate (R) | 6,239 | -0.0002 | -6.39 | < .001 | -0.06 | [-.07, -.05] |
| Posterior Cingulate (L) | 6,250 | -0.0002 | -6.39 | < .001 | -0.06 | [-.07, -.05] |
| Supramarginal (L) | 6,130 | -0.0002 | -6.25 | < .001 | -0.06 | [-.07, -.05] |
| Banks of the Superior Temporal Sulcus (R) | 6,308 | -0.0002 | -6.08 | < .001 | -0.05 | [-.06, -.05] |
| Fusiform (L) | 6,251 | -0.0002 | -5.95 | < .001 | -0.05 | [-.06, -.04] |
| Superior Parietal (R) | 6,236 | -0.0002 | -5.94 | < .001 | -0.05 | [-.06, -.04] |
| Inferior Temporal (R) | 6,224 | -0.0002 | -5.88 | < .001 | -0.05 | [-.06, -.04] |
| Parahippocampal (L) | 6,295 | -0.0003 | -5.78 | < .001 | -0.05 | [-.06, -.04] |
| Insula (L) | 6,273 | -0.0003 | -5.75 | < .001 | -0.05 | [-.06, -.04] |
| Fusiform (R) | 6,259 | -0.0002 | -5.70 | < .001 | -0.05 | [-.06, -.04] |
| Insula (R) | 6,271 | -0.0003 | -5.54 | < .001 | -0.05 | [-.06, -.04] |
| Supramarginal (R) | 6,148 | -0.0002 | -5.47 | < .001 | -0.05 | [-.06, -.04] |
| Parahippocampal (R) | 6,287 | -0.0003 | -5.34 | < .001 | -0.05 | [-.06, -.04] |
| Pars Opercularis (R) | 6,255 | -0.0002 | -5.34 | < .001 | -0.05 | [-.06, -.04] |
| Superior Parietal (L) | 6,256 | -0.0002 | -5.08 | < .001 | -0.05 | [-.05, -.04] |
| Banks of the Superior Temporal Sulcus (L) | 6,282 | -0.0002 | -4.75 | < .001 | -0.04 | [-.05, -.03] |
| Rostral Anterior Cingulate (R) | 6,276 | -0.0003 | -4.59 | < .001 | -0.04 | [-.05, -.03] |
| Rostral Middle Frontal (R) | 6,239 | -0.0002 | -4.49 | < .001 | -0.04 | [-.05, -.03] |
| Rostral Anterior Cingulate (L) | 6,283 | -0.0003 | -4.42 | < .001 | -0.04 | [-.05, -.03] |
| Middle Temporal (R) | 6,149 | -0.0002 | -4.36 | < .001 | -0.04 | [-.05, -.03] |
| Lingual (R) | 6,284 | -0.0001 | -4.12 | < .001 | -0.04 | [-.05, -.03] |
| Caudal Anterior Cingulate (L) | 6,277 | -0.0002 | -4.09 | < .001 | -0.04 | [-.05, -.03] |
| Lingual (L) | 6,282 | -0.0001 | -3.85 | < .001 | -0.03 | [-.04, -.03] |
| Pars Orbitalis (R) | 6,265 | -0.0002 | -3.78 | < .001 | -0.03 | [-.04, -.02] |
| Rostral Middle Frontal (L) | 6,225 | -0.0001 | -3.78 | < .001 | -0.03 | [-.04, -.02] |
| Inferior Temporal (L) | 6,230 | -0.0001 | -3.74 | < .001 | -0.03 | [-.04, -.02] |
| Paracentral (R) | 6,260 | -0.0001 | -3.66 | < .001 | -0.03 | [-.04, -.02] |
| Paracentral (L) | 6,254 | -0.0001 | -3.64 | < .001 | -0.03 | [-.04, -.02] |
| Pars Triangularis (R) | 6,241 | -0.0001 | -3.57 | < .001 | -0.03 | [-.04, -.02] |
| Medial Orbital Frontal (L) | 6,259 | -0.0002 | -3.55 | < .001 | -0.03 | [-.04, -.02] |
| Transverse Temporal (L) | 6,288 | -0.0002 | -3.35 | .001 | -0.03 | [-.04, -.02] |
| Pars Opercularis (L) | 6,233 | -0.0001 | -3.31 | .001 | -0.03 | [-.04, -.02] |
| Medial Orbital Frontal (R) | 6,262 | -0.0001 | -3.13 | .002 | -0.03 | [-.04, -.02] |
| Postcentral (R) | 6,151 | -0.0001 | -3.07 | .002 | -0.03 | [-.04, -.02] |
| Caudal Middle Frontal (L) | 6,217 | -0.0001 | -2.92 | .003 | -0.03 | [-.04, -.02] |
| Caudal Anterior Cingulate (R) | 6,226 | -0.0001 | -2.91 | .004 | -0.03 | [-.04, -.02] |
| Transverse Temporal (R) | 6,276 | -0.0002 | -2.88 | .004 | -0.03 | [-.03, -.02] |
| Middle Temporal (L) | 6,169 | -0.0001 | -2.86 | .004 | -0.03 | [-.03, -.02] |
| Superior Frontal (R) | 6,259 | -0.0001 | -2.85 | .004 | -0.03 | [-.03, -.02] |
| Superior Frontal (L) | 6,245 | -0.0001 | -2.83 | .005 | -0.03 | [-.03, -.02] |
| Superior Temporal (R) | 6,272 | -0.0001 | -2.83 | .005 | -0.03 | [-.03, -.02] |
| Lateral Orbital Frontal (R) | 6,264 | -0.0001 | -2.71 | .007 | -0.02 | [-.03, -.02] |
| Postcentral (L) | 6,237 | -0.0001 | -2.69 | .007 | -0.02 | [-.03, -.02] |
| Lateral Orbital Frontal (L) | 6,258 | -0.0001 | -2.57 | .010 | -0.02 | [-.03, -.01] |
| Cuneus (L) | 6,283 | -0.0001 | -2.57 | .010 | -0.02 | [-.03, -.01] |
| Pars Triangularis (L) | 6,233 | -0.0001 | -2.50 | .013 | -0.02 | [-.03, -.01] |
| Lateral Occipital (R) | 6,291 | -0.0001 | -2.48 | .013 | -0.02 | [-.03, -.01] |
| Pars Orbitalis (L) | 6,244 | -0.0001 | -2.44 | .015 | -0.02 | [-.03, -.01] |
| Cuneus (R) | 6,287 | -0.0001 | -2.38 | .017 | -0.02 | [-.03, -.01] |
| Superior Temporal (L) | 6,265 | -0.0001 | -2.27 | .023 | -0.02 | [-.03, -.01] |
| Caudal Middle Frontal (R) | 6,209 | -0.0001 | -2.24 | .025 | -0.02 | [-.03, -.01] |
| Temporal Pole (R) | 6,156 | -0.0002 | -1.81 | .070 | -0.02 | [-.03, -.01] |
| Entorhinal (R) | 6,248 | -0.0002 | -1.38 | .167 | -0.01 | [-.02, -.003] |
| Precentral (R) | 6,022 | 0.0000 | -1.25 | .213 | -0.01 | [-.02, -.002] |
| Lateral Occipital (L) | 6,296 | 0.0000 | -1.06 | .289 | -0.01 | [-.02, -.001] |
| Pericalcarine (L) | 6,302 | 0.0000 | -0.94 | .346 | -0.01 | [-.02, .001] |
| Frontal Pole (R) | 6,239 | -0.0001 | -0.87 | .387 | -0.01 | [-.02, .001] |
| Temporal Pole (L) | 6,192 | 0.0001 | 0.80 | .426 | 0.01 | [-.002, .02] |
| Pericalcarine (R) | 6,295 | 0.0000 | 0.78 | .438 | 0.01 | [-.002, .02] |
| Frontal Pole (L) | 6,225 | -0.0001 | -0.71 | .478 | -0.01 | [-.02, .003] |
| Precentral (L) | 6,128 | 0.0000 | -0.45 | .654 | 0.00 | [-.01, .005] |
| Entorhinal (L) | 6,227 | 0.0000 | -0.33 | .741 | 0.00 | [-.01, .01] |

*Note*. Cortical regions are sorted vertically by *p*-value (uncorrected) for Sex × Age associations. Analyses included fixed effects of sex, age, age^2^, Sex × Age, Sex × Age^2^, and whole-brain volume. The random-effects structure included random intercepts of participant ID and magnetic resonance imaging (MRI) scanner serial number. Associations that passed false-discovery-rate (FDR) correction are shaded. *n* = number of participants included in analysis of that region. *t* = *t*-statistic. *b* = unstandardized regression coefficient [i.e., difference in change in cortical thickness (mm) (relative to the coefficients in Supplementary Table 1) with a one-month increase in age in participants assigned female sex at birth]. 95% CI = 95% confidence interval. *r_p_* = partial correlation coefficient (i.e., effect size).

**Extended Data Figure 3-2.** Sex × Age (in months) interactions on sulcal depth (mm).

| **Cortical Region (Hemisphere)** | ***n*** | ***b*** | ***t*** | ***p*** | ***r_p_*** | **95% CI (*r_p_*)** |
| --- | --- | --- | --- | --- | --- | --- |
| Lingual (R) | 6,272 | 0.0002 | 4.38 | < .001 | 0.04 | [.03, .05] |
| Temporal Pole (R) | 6,264 | 0.0008 | 3.91 | < .001 | 0.03 | [.03, .04] |
| Caudal Anterior Cingulate (R) | 6,262 | 0.0003 | 3.61 | < .001 | 0.03 | [.02, .04] |
| Superior Parietal (R) | 6,269 | -0.0002 | -3.54 | < .001 | -0.03 | [-.04, -.02] |
| Fusiform (L) | 6,292 | 0.0002 | 3.02 | .003 | 0.03 | [.02, .04] |
| Rostral Middle Frontal (L) | 6,282 | -0.0001 | -2.99 | .003 | -0.03 | [-.04, -.02] |
| Fusiform (R) | 6,292 | 0.0002 | 2.85 | .004 | 0.03 | [.02, .03] |
| Superior Parietal (L) | 6,289 | -0.0001 | -2.83 | .005 | -0.03 | [-.03, -.02] |
| Parahippocampal (R) | 6,294 | -0.0003 | -2.75 | .006 | -0.02 | [-.03, -.02] |
| Rostral Anterior Cingulate (R) | 6,282 | 0.0003 | 2.74 | .006 | 0.02 | [.02, .03] |
| Rostral Middle Frontal (R) | 6,292 | -0.0001 | -2.59 | .010 | -0.02 | [-.03, -.01] |
| Lingual (L) | 6,271 | 0.0001 | 2.48 | .013 | 0.02 | [.01, .03] |
| Pars Triangularis (R) | 6,260 | -0.0002 | -2.41 | .016 | -0.02 | [-.03, -.01] |
| Precentral (R) | 6,235 | 0.0001 | 2.35 | .019 | 0.02 | [.01, .03] |
| Inferior Temporal (L) | 6,290 | -0.0001 | -2.16 | .031 | -0.02 | [-.03, -.01] |
| Caudal Anterior Cingulate (L) | 6,270 | 0.0002 | 2.08 | .038 | 0.02 | [.01, .03] |
| Medial Orbital Frontal (L) | 6,294 | 0.0002 | 2.00 | .045 | 0.02 | [.01, .03] |
| Inferior Parietal (L) | 6,283 | -0.0001 | -1.89 | .059 | -0.02 | [-.03, -.01] |
| Precuneus (R) | 6,291 | 0.0001 | 1.86 | .062 | 0.02 | [.01, .03] |
| Lateral Orbital Frontal (L) | 6,284 | -0.0001 | -1.81 | .070 | -0.02 | [-.03, -.01] |
| Transverse Temporal (L) | 6,306 | -0.0002 | -1.77 | .077 | -0.02 | [-.02, -.01] |
| Superior Temporal (R) | 6,278 | -0.0001 | -1.76 | .078 | -0.02 | [-.02, -.01] |
| Isthmus Cingulate (R) | 6,254 | 0.0001 | 1.74 | .081 | 0.02 | [.01, .02] |
| Transverse Temporal (R) | 6,295 | -0.0002 | -1.69 | .090 | -0.02 | [-.02, -.01] |
| Banks of the Superior Temporal Sulcus (L) | 6,150 | -0.0002 | -1.60 | .109 | -0.01 | [-.02, -.01] |
| Isthmus Cingulate (L) | 6,252 | 0.0001 | 1.49 | .135 | 0.01 | [.004, .02] |
| Parahippocampal (L) | 6,310 | -0.0002 | -1.41 | .158 | -0.01 | [-.02, -.004] |
| Superior Frontal (L) | 6,294 | 0.0000 | 1.36 | .173 | 0.01 | [.003, .02] |
| Insula (L) | 6,311 | -0.0002 | -1.32 | .186 | -0.01 | [-.02, -.003] |
| Paracentral (L) | 6,257 | -0.0001 | -1.28 | .201 | -0.01 | [-.02, -.002] |
| Entorhinal (R) | 6,310 | 0.0003 | 1.25 | .211 | 0.01 | [.002, .02] |
| Banks of the Superior Temporal Sulcus (R) | 6,111 | -0.0001 | -1.14 | .255 | -0.01 | [-.02, -.001] |
| Cuneus (R) | 6,296 | -0.0001 | -1.12 | .263 | -0.01 | [-.02, -.001] |
| Paracentral (R) | 6,267 | -0.0001 | -1.09 | .278 | -0.01 | [-.02, -.001] |
| Frontal Pole (R) | 6,264 | -0.0002 | -1.08 | .279 | -0.01 | [-.02, -.001] |
| Supramarginal (L) | 6,264 | 0.0001 | 1.07 | .284 | 0.01 | [.001, .02] |
| Rostral Anterior Cingulate (L) | 6,298 | 0.0001 | 1.06 | .291 | 0.01 | [.0005, .02] |
| Lateral Occipital (R) | 6,302 | 0.0000 | -0.98 | .329 | -0.01 | [-.02, .0002] |
| Pars Orbitalis (L) | 6,285 | -0.0001 | -0.96 | .337 | -0.01 | [-.02, .0004] |
| Temporal Pole (L) | 6,260 | 0.0002 | 0.96 | .339 | 0.01 | [-.0004, .02] |
| Caudal Middle Frontal (L) | 6,228 | -0.0001 | -0.94 | .348 | -0.01 | [-.02, .001] |
| Precentral (L) | 6,218 | 0.0000 | 0.92 | .359 | 0.01 | [-.001, .02] |
| Middle Temporal (L) | 6,276 | 0.0001 | 0.91 | .361 | 0.01 | [-.001, .02] |
| Precuneus (L) | 6,299 | 0.0000 | 0.89 | .376 | 0.01 | [-.001, .02] |
| Entorhinal (L) | 6,312 | 0.0002 | 0.73 | .468 | 0.01 | [-.002, .02] |
| Superior Temporal (L) | 6,266 | 0.0000 | -0.71 | .480 | -0.01 | [-.02, .003] |
| Posterior Cingulate (L) | 6,278 | 0.0000 | 0.67 | .503 | 0.01 | [-.003, .01] |
| Inferior Parietal (R) | 6,291 | 0.0000 | -0.67 | .503 | -0.01 | [-.01, .003] |
| Pars Triangularis (L) | 6,253 | 0.0001 | 0.62 | .538 | 0.01 | [-.003, .01] |
| Cuneus (L) | 6,290 | 0.0001 | 0.60 | .550 | 0.01 | [-.004, .01] |
| Pars Opercularis (R) | 6,169 | 0.0001 | 0.56 | .577 | 0.01 | [-.004, .01] |
| Pars Opercularis (L) | 6,225 | 0.0000 | -0.55 | .585 | 0.00 | [-.01, .004] |
| Insula (R) | 6,278 | -0.0001 | -0.51 | .609 | 0.00 | [-.01, .004] |
| Posterior Cingulate (R) | 6,287 | 0.0000 | 0.49 | .625 | 0.00 | [-.005, .01] |
| Postcentral (R) | 6,178 | 0.0000 | -0.41 | .682 | 0.00 | [-.01, .01] |
| Lateral Occipital (L) | 6,304 | 0.0000 | 0.38 | .704 | 0.00 | [-.01, .01] |
| Postcentral (L) | 6,252 | 0.0000 | 0.37 | .711 | 0.00 | [-.01, .01] |
| Pars Orbitalis (R) | 6,284 | 0.0000 | -0.36 | .718 | 0.00 | [-.01, .01] |
| Middle Temporal (R) | 6,287 | 0.0000 | 0.32 | .751 | 0.00 | [-.01, .01] |
| Caudal Middle Frontal (R) | 6,217 | 0.0000 | 0.23 | .819 | 0.00 | [-.01, .01] |
| Lateral Orbital Frontal (R) | 6,283 | 0.0000 | -0.18 | .859 | 0.00 | [-.01, .01] |
| Superior Frontal (R) | 6,293 | 0.0000 | -0.16 | .870 | 0.00 | [-.01, .01] |
| Inferior Temporal (R) | 6,285 | 0.0000 | 0.15 | .878 | 0.00 | [-.01, .01] |
| Pericalcarine (R) | 6,227 | 0.0000 | -0.15 | .884 | 0.00 | [-.01, .01] |
| Medial Orbital Frontal (R) | 6,278 | 0.0000 | -0.14 | .891 | 0.00 | [-.01, .01] |
| Frontal Pole (L) | 6,238 | 0.0000 | 0.13 | .900 | 0.00 | [-.01, .01] |
| Supramarginal (R) | 6,269 | 0.0000 | 0.09 | .927 | 0.00 | [-.01, .01] |
| Pericalcarine (L) | 6,219 | 0.0000 | 0.08 | .937 | 0.00 | [-.01, .01] |

*Note*. Cortical regions are sorted vertically by *p*-value (uncorrected) for Sex × Age associations. Analyses included fixed effects of sex, age, age^2^, Sex × Age, Sex × Age^2^, and whole-brain volume. The random-effects structure included random intercepts of participant ID and magnetic resonance imaging (MRI) scanner serial number. Associations that passed false-discovery-rate (FDR) correction are shaded. *n* = number of participants included in analysis of that region. *t* = *t*-statistic. *b* = unstandardized regression coefficient [i.e., difference in change in cortical sulcal depth (mm) (relative to the coefficients in Supplementary Table 2) with a one-month increase in age in participants assigned female sex at birth]. 95% CI = 95% confidence interval. *r_p_* = partial correlation coefficient (i.e., effect size).

**Extended Data Figure 3-3.** Sex × Age (in months) interactions on cortical surface area (mm^2^).

| **Cortical Region (Hemisphere)** | ***n*** | ***b*** | ***t*** | ***p*** | ***r_p_*** | **95% CI (*r_p_*)** |
| --- | --- | --- | --- | --- | --- | --- |
| Rostral Middle Frontal (L) | 6,308 | -0.4895 | -4.04 | < .001 | -0.04 | [-.04, -.03] |
| Banks of the Superior Temporal Sulcus (L) | 6,245 | -0.1166 | -4.01 | < .001 | -0.04 | [-.04, -.03] |
| Superior Parietal (R) | 6,267 | 0.4306 | 3.18 | .001 | 0.03 | [.02, .04] |
| Rostral Middle Frontal (R) | 6,308 | -0.4718 | -3.18 | .001 | -0.03 | [-.04, -.02] |
| Caudal Anterior Cingulate (R) | 6,256 | -0.0623 | -2.84 | .005 | -0.03 | [-.03, -.02] |
| Postcentral (R) | 6,237 | 0.2192 | 2.80 | .005 | 0.03 | [.02, .03] |
| Caudal Middle Frontal (R) | 6,277 | -0.2139 | -2.79 | .005 | -0.02 | [-.03, -.02] |
| Superior Frontal (R) | 6,277 | -0.4369 | -2.79 | .005 | -0.02 | [-.03, -.02] |
| Precuneus (R) | 6,271 | 0.1855 | 2.76 | .006 | 0.02 | [.02, .03] |
| Pars Triangularis (L) | 6,298 | -0.0822 | -2.73 | .006 | -0.02 | [-.03, -.02] |
| Temporal Pole (R) | 6,243 | 0.0463 | 2.62 | .009 | 0.02 | [.01, .03] |
| Cuneus (L) | 6,289 | 0.0669 | 2.61 | .009 | 0.02 | [.01, .03] |
| Precentral (R) | 6,243 | -0.2307 | -2.43 | .015 | -0.02 | [-.03, -.01] |
| Transverse Temporal (R) | 6,232 | -0.0186 | -2.23 | .026 | -0.02 | [-.03, -.01] |
| Pars Opercularis (L) | 6,226 | -0.0779 | -2.12 | .034 | -0.02 | [-.03, -.01] |
| Superior Temporal (L) | 6,266 | -0.1277 | -2.12 | .034 | -0.02 | [-.03, -.01] |
| Postcentral (L) | 6,239 | 0.1419 | 2.07 | .038 | 0.02 | [.01, .03] |
| Paracentral (L) | 6,256 | 0.0512 | 1.99 | .047 | 0.02 | [.01, .03] |
| Supramarginal (L) | 6,276 | 0.2049 | 1.96 | .050 | 0.02 | [.01, .03] |
| Parahippocampal (R) | 6,222 | 0.0345 | 1.95 | .051 | 0.02 | [.01, .03] |
| Caudal Middle Frontal (L) | 6,283 | -0.1142 | -1.88 | .060 | -0.02 | [-.03, -.01] |
| Parahippocampal (L) | 6,196 | 0.0322 | 1.75 | .080 | 0.02 | [.01, .02] |
| Superior Frontal (L) | 6,286 | -0.2062 | -1.71 | .087 | -0.02 | [-.02, -.01] |
| Inferior Parietal (R) | 6,299 | 0.1691 | 1.71 | .088 | 0.02 | [.01, .02] |
| Supramarginal (R) | 6,266 | 0.1770 | 1.61 | .108 | 0.01 | [.01, .02] |
| Middle Temporal (R) | 6,289 | -0.0879 | -1.58 | .114 | -0.01 | [-.02, -.01] |
| Temporal Pole (L) | 6,274 | 0.0293 | 1.58 | .115 | 0.01 | [.01, .02] |
| Banks of the Superior Temporal Sulcus (R) | 6,267 | -0.0347 | -1.54 | .123 | -0.01 | [-.02, -.005] |
| Inferior Temporal (L) | 6,317 | 0.0830 | 1.49 | .137 | 0.01 | [.004, .02] |
| Lingual (L) | 6,289 | 0.0621 | 1.45 | .147 | 0.01 | [.004, .02] |
| Insula (L) | 6,266 | 0.1078 | 1.42 | .155 | 0.01 | [.004, .02] |
| Rostral Anterior Cingulate (L) | 6,301 | 0.0421 | 1.37 | .171 | 0.01 | [.003, .02] |
| Superior Parietal (L) | 6,261 | 0.1611 | 1.33 | .183 | 0.01 | [.003, .02] |
| Middle Temporal (L) | 6,291 | -0.0766 | -1.31 | .189 | -0.01 | [-.02, -.003] |
| Pars Triangularis (R) | 6,269 | -0.0530 | -1.24 | .216 | -0.01 | [-.02, -.002] |
| Frontal Pole (R) | 6,277 | -0.0169 | -1.22 | .223 | -0.01 | [-.02, -.002] |
| Paracentral (R) | 6,242 | 0.0351 | 1.17 | .240 | 0.01 | [.002, .02] |
| Pericalcarine (L) | 6,307 | 0.0343 | 1.15 | .250 | 0.01 | [.001, .02] |
| Precentral (L) | 6,250 | -0.0922 | -1.15 | .251 | -0.01 | [-.02, -.001] |
| Medial Orbital Frontal (R) | 6,290 | 0.0620 | 1.11 | .267 | 0.01 | [.001, .02] |
| Isthmus Cingulate (L) | 6,222 | 0.0258 | 1.09 | .276 | 0.01 | [.001, .02] |
| Pars Orbitalis (L) | 6,312 | 0.0162 | 1.04 | .298 | 0.01 | [.0004, .02] |
| Lingual (R) | 6,296 | 0.0413 | 0.91 | .365 | 0.01 | [-.001, .02] |
| Precuneus (L) | 6,282 | 0.0547 | 0.89 | .374 | 0.01 | [-.001, .02] |
| Posterior Cingulate (R) | 6,265 | -0.0205 | -0.87 | .385 | -0.01 | [-.02, .001] |
| Lateral Orbital Frontal (L) | 6,305 | 0.0438 | 0.74 | .457 | 0.01 | [-.002, .02] |
| Fusiform (L) | 6,298 | -0.0314 | -0.74 | .459 | -0.01 | [-.02, .002] |
| Entorhinal (L) | 6,246 | -0.0173 | -0.74 | .462 | -0.01 | [-.02, .002] |
| Pars Orbitalis (R) | 6,307 | -0.0129 | -0.71 | .475 | -0.01 | [-.02, .003] |
| Posterior Cingulate (L) | 6,251 | -0.0156 | -0.70 | .487 | -0.01 | [-.02, .003] |
| Inferior Temporal (R) | 6,313 | 0.0281 | 0.55 | .583 | 0.00 | [-.004, .01] |
| Entorhinal (R) | 6,225 | 0.0111 | 0.54 | .587 | 0.00 | [-.004, .01] |
| Lateral Occipital (L) | 6,313 | 0.0336 | 0.53 | .597 | 0.00 | [-.004, .01] |
| Medial Orbital Frontal (L) | 6,276 | 0.0305 | 0.45 | .649 | 0.00 | [-.005, .01] |
| Rostral Anterior Cingulate (R) | 6,313 | -0.0099 | -0.45 | .656 | 0.00 | [-.01, .005] |
| Superior Temporal (R) | 6,255 | -0.0223 | -0.43 | .664 | 0.00 | [-.01, .01] |
| Lateral Occipital (R) | 6,308 | -0.0269 | -0.40 | .692 | 0.00 | [-.01, .01] |
| Frontal Pole (L) | 6,289 | -0.0042 | -0.39 | .699 | 0.00 | [-.01, .01] |
| Insula (R) | 6,267 | -0.0257 | -0.32 | .747 | 0.00 | [-.01, .01] |
| Transverse Temporal (L) | 6,209 | 0.0030 | 0.28 | .777 | 0.00 | [-.01, .01] |
| Cuneus (R) | 6,284 | 0.0060 | 0.21 | .837 | 0.00 | [-.01, .01] |
| Pericalcarine (R) | 6,309 | -0.0054 | -0.17 | .867 | 0.00 | [-.01, .01] |
| Fusiform (R) | 6,305 | 0.0063 | 0.15 | .878 | 0.00 | [-.01, .01] |
| Inferior Parietal (L) | 6,296 | -0.0118 | -0.15 | .879 | 0.00 | [-.01, .01] |
| Caudal Anterior Cingulate (L) | 6,201 | -0.0023 | -0.13 | .899 | 0.00 | [-.01, .01] |
| Lateral Orbital Frontal (R) | 6,291 | 0.0091 | 0.10 | .921 | 0.00 | [-.01, .01] |
| Isthmus Cingulate (R) | 6,235 | 0.0019 | 0.08 | .936 | 0.00 | [-.01, .01] |
| Pars Opercularis (R) | 6,234 | 0.0027 | 0.07 | .943 | 0.00 | [-.01, .01] |

*Note*. Cortical regions are sorted vertically by *p*-value (uncorrected) for Sex × Age associations. Analyses included fixed effects of sex, age, age^2^, Sex × Age, Sex × Age^2^, and whole-brain volume. The random-effects structure included random intercepts of participant ID and magnetic resonance imaging (MRI) scanner serial number. Associations that passed false-discovery-rate (FDR) correction are shaded. *n* = number of participants included in analysis of that region. *t* = *t*-statistic. *b* = unstandardized regression coefficient [i.e., difference in change in cortical surface area (mm^2^) (relative to the coefficients in Supplementary Table 3) with a one-month increase in age in participants assigned female sex at birth]. 95% CI = 95% confidence interval. *r_p_* = partial correlation coefficient (i.e., effect size).

**Extended Data Figure 3-4.** Sex × Age (in months) interactions on cortical volume (mm^3^).

| **Cortical Region (Hemisphere)** | ***n*** | ***b*** | ***t*** | ***p*** | ***r_p_*** | **95% CI (*r_p_*)** |
| --- | --- | --- | --- | --- | --- | --- |
| Inferior Parietal (L) | 6,297 | -1.3690 | -5.60 | < .001 | -0.05 | [-.06, -.04] |
| Insula (R) | 6,267 | -0.9771 | -4.58 | < .001 | -0.04 | [-.05, -.03] |
| Isthmus Cingulate (R) | 6,238 | -0.2998 | -4.41 | < .001 | -0.04 | [-.05, -.03] |
| Banks of the Superior Temporal Sulcus (L) | 6,225 | -0.3875 | -4.12 | < .001 | -0.04 | [-.05, -.03] |
| Pars Opercularis (L) | 6,227 | -0.5060 | -4.09 | < .001 | -0.04 | [-.05, -.03] |
| Rostral Middle Frontal (L) | 6,300 | -1.5854 | -4.00 | < .001 | -0.04 | [-.04, -.03] |
| Inferior Temporal (R) | 6,308 | -0.8298 | -3.98 | < .001 | -0.04 | [-.04, -.03] |
| Middle Temporal (R) | 6,303 | -0.8133 | -3.92 | < .001 | -0.03 | [-.04, -.03] |
| Posterior Cingulate (R) | 6,267 | -0.2638 | -3.86 | < .001 | -0.03 | [-.04, -.03] |
| Superior Frontal (R) | 6,288 | -1.9561 | -3.83 | < .001 | -0.03 | [-.04, -.03] |
| Caudal Middle Frontal (L) | 6,294 | -0.7448 | -3.71 | < .001 | -0.03 | [-.04, -.02] |
| Rostral Middle Frontal (R) | 6,303 | -1.7394 | -3.71 | < .001 | -0.03 | [-.04, -.02] |
| Fusiform (L) | 6,288 | -0.6560 | -3.53 | < .001 | -0.03 | [-.04, -.02] |
| Inferior Parietal (R) | 6,293 | -1.0517 | -3.38 | .001 | -0.03 | [-.04, -.02] |
| Pars Orbitalis (R) | 6,302 | -0.2473 | -3.32 | .001 | -0.03 | [-.04, -.02] |
| Pars Triangularis (L) | 6,279 | -0.3514 | -3.31 | .001 | -0.03 | [-.04, -.02] |
| Fusiform (R) | 6,295 | -0.5596 | -3.20 | .001 | -0.03 | [-.04, -.02] |
| Caudal Anterior Cingulate (R) | 6,241 | -0.2238 | -3.15 | .002 | -0.03 | [-.04, -.02] |
| Caudal Middle Frontal (R) | 6,284 | -0.7750 | -3.11 | .002 | -0.03 | [-.04, -.02] |
| Rostral Anterior Cingulate (R) | 6,313 | -0.2433 | -3.00 | .003 | -0.03 | [-.04, -.02] |
| Banks of the Superior Temporal Sulcus (R) | 6,254 | -0.2131 | -2.92 | .004 | -0.03 | [-.04, -.02] |
| Precuneus (L) | 6,297 | -0.5757 | -2.89 | .004 | -0.03 | [-.03, -.02] |
| Middle Temporal (L) | 6,296 | -0.6032 | -2.85 | .004 | -0.03 | [-.03, -.02] |
| Isthmus Cingulate (L) | 6,239 | -0.1807 | -2.69 | .007 | -0.02 | [-.03, -.02] |
| Pars Opercularis (R) | 6,232 | -0.3611 | -2.69 | .007 | -0.02 | [-.03, -.02] |
| Superior Temporal (L) | 6,282 | -0.5441 | -2.69 | .007 | -0.02 | [-.03, -.02] |
| Pericalcarine (R) | 6,287 | 0.1941 | 2.55 | .011 | 0.02 | [.01, .03] |
| Lingual (R) | 6,291 | -0.3150 | -2.45 | .014 | -0.02 | [-.03, -.01] |
| Posterior Cingulate (L) | 6,265 | -0.1625 | -2.37 | .018 | -0.02 | [-.03, -.01] |
| Lingual (L) | 6,288 | -0.2838 | -2.34 | .019 | -0.02 | [-.03, -.01] |
| Insula (L) | 6,272 | -0.4585 | -2.29 | .022 | -0.02 | [-.03, -.01] |
| Rostral Anterior Cingulate (L) | 6,298 | -0.2271 | -2.15 | .031 | -0.02 | [-.03, -.01] |
| Superior Frontal (L) | 6,296 | -0.8219 | -1.99 | .047 | -0.02 | [-.03, -.01] |
| Transverse Temporal (R) | 6,207 | -0.0570 | -1.81 | .070 | -0.02 | [-.03, -.01] |
| Inferior Temporal (L) | 6,309 | -0.3826 | -1.69 | .090 | -0.02 | [-.02, -.01] |
| Pericalcarine (L) | 6,296 | 0.1048 | 1.60 | .110 | 0.01 | [.01, .02] |
| Precentral (L) | 6,276 | -0.3698 | -1.55 | .122 | -0.01 | [-.02, -.005] |
| Cuneus (L) | 6,256 | 0.1197 | 1.54 | .123 | 0.01 | [.005, .02] |
| Temporal Pole (L) | 6,263 | 0.1927 | 1.52 | .128 | 0.01 | [.005, .02] |
| Precuneus (R) | 6,272 | -0.3201 | -1.48 | .139 | -0.01 | [-.02, -.004] |
| Superior Parietal (L) | 6,270 | -0.5567 | -1.47 | .143 | -0.01 | [-.02, -.004] |
| Lateral Orbital Frontal (R) | 6,277 | -0.3248 | -1.45 | .148 | -0.01 | [-.02, -.004] |
| Parahippocampal (L) | 6,221 | -0.1007 | -1.44 | .150 | -0.01 | [-.02, -.004] |
| Lateral Orbital Frontal (L) | 6,293 | -0.2047 | -1.24 | .216 | -0.01 | [-.02, -.002] |
| Postcentral (L) | 6,263 | 0.2479 | 1.22 | .222 | 0.01 | [.002, .02] |
| Supramarginal (L) | 6,287 | -0.4044 | -1.21 | .228 | -0.01 | [-.02, -.002] |
| Lateral Occipital (L) | 6,300 | 0.2401 | 1.11 | .267 | 0.01 | [.001, .02] |
| Parahippocampal (R) | 6,229 | -0.0672 | -1.02 | .308 | -0.01 | [-.02, -.0002] |
| Medial Orbital Frontal (L) | 6,280 | -0.1869 | -1.00 | .318 | -0.01 | [-.02, .00001] |
| Caudal Anterior Cingulate (L) | 6,209 | -0.0646 | -1.00 | .319 | -0.01 | [-.02, .00002] |
| Temporal Pole (R) | 6,245 | -0.1065 | -0.88 | .378 | -0.01 | [-.02, .001] |
| Frontal Pole (R) | 6,246 | -0.0597 | -0.83 | .406 | -0.01 | [-.02, .002] |
| Cuneus (R) | 6,249 | 0.0703 | 0.72 | .471 | 0.01 | [-.003, .02] |
| Pars Triangularis (R) | 6,262 | -0.1077 | -0.72 | .474 | -0.01 | [-.02, .003] |
| Postcentral (R) | 6,276 | 0.1452 | 0.69 | .487 | 0.01 | [-.003, .02] |
| Pars Orbitalis (L) | 6,300 | -0.0423 | -0.63 | .530 | -0.01 | [-.01, .003] |
| Paracentral (L) | 6,243 | -0.0589 | -0.62 | .538 | -0.01 | [-.01, .003] |
| Precentral (R) | 6,229 | -0.1662 | -0.58 | .559 | -0.01 | [-.01, .004] |
| Frontal Pole (L) | 6,262 | -0.0268 | -0.47 | .641 | 0.00 | [-.01, .005] |
| Superior Temporal (R) | 6,270 | -0.0805 | -0.46 | .648 | 0.00 | [-.01, .005] |
| Superior Parietal (R) | 6,264 | 0.1895 | 0.44 | .660 | 0.00 | [-.005, .01] |
| Paracentral (R) | 6,243 | -0.0442 | -0.40 | .688 | 0.00 | [-.01, .01] |
| Medial Orbital Frontal (R) | 6,280 | -0.0527 | -0.30 | .762 | 0.00 | [-.01, .01] |
| Supramarginal (R) | 6,282 | 0.0862 | 0.24 | .808 | 0.00 | [-.01, .01] |
| Lateral Occipital (R) | 6,297 | -0.0437 | -0.19 | .848 | 0.00 | [-.01, .01] |
| Transverse Temporal (L) | 6,256 | -0.0043 | -0.11 | .910 | 0.00 | [-.01, .01] |
| Entorhinal (R) | 6,163 | -0.0086 | -0.08 | .935 | 0.00 | [-.01, .01] |
| Entorhinal (L) | 6,139 | -0.0058 | -0.05 | .956 | 0.00 | [-.01, .01] |

*Note*. Cortical regions are sorted vertically by *p*-value (uncorrected) for Sex × Age associations. Analyses included fixed effects of sex, age, age^2^, Sex × Age, Sex × Age^2^, and whole-brain volume. The random-effects structure included random intercepts of participant ID and magnetic resonance imaging (MRI) scanner serial number. Associations that passed false-discovery-rate (FDR) correction are shaded. *n* = number of participants included in analysis of that region. *t* = *t*-statistic. *b* = unstandardized regression coefficient [i.e., difference in change in cortical volume (mm^3^) (relative to the coefficients in Supplementary Table 4) with a one-month increase in age in participants assigned female sex at birth]. 95% CI = 95% confidence interval. *r_p_* = partial correlation coefficient (i.e., effect size).

**Extended Data Figure 7-1.** Main effects of sex-at-birth within APC_CT_*APC_CSA_ circular analyses.

| **Cortical Region (Hemisphere)** | ***SS_B_*** | ***SS_W_*** | ***F*** | ***p*** | **% Variance** |
| --- | --- | --- | --- | --- | --- |
| Caudal Middle Frontal (L) | 39.0 | 3710.4 | 91.52 | < .001 | 1.04 |
| Middle Temporal (R) | 24.6 | 2607.7 | 62.88 | < .001 | 0.93 |
| Middle Temporal (L) | 24.8 | 2672.5 | 62.67 | < .001 | 0.92 |
| Caudal Middle Frontal (R) | 31.6 | 3498.3 | 72.85 | < .001 | 0.9 |
| Superior Frontal (R) | 21.6 | 2542 | 55.84 | < .001 | 0.84 |
| Superior Temporal (L) | 22.6 | 3154.7 | 52.79 | < .001 | 0.71 |
| Rostral Middle Frontal (L) | 18.7 | 2657.3 | 47.41 | < .001 | 0.7 |
| Precentral (R) | 22.1 | 3538.9 | 51.67 | < .001 | 0.62 |
| Rostral Middle Frontal (R) | 14.7 | 2431.5 | 39.4 | < .001 | 0.6 |
| Pars Triangularis (R) | 16.2 | 3003.4 | 38.78 | < .001 | 0.54 |
| Pars Opercularis (L) | 18.2 | 3552.8 | 41.85 | < .001 | 0.51 |
| Inferior Temporal (R) | 14.0 | 2951.9 | 33.79 | < .001 | 0.47 |
| Inferior Temporal (L) | 13.0 | 2839 | 32.05 | < .001 | 0.46 |
| Frontal Pole (L) | 15.3 | 3646.7 | 35.16 | < .001 | 0.42 |
| Pars Opercularis (R) | 15.1 | 3616.6 | 34.74 | < .001 | 0.42 |
| Banks of the Superior Temporal Sulcus (L) | 12.5 | 3036.7 | 29.75 | < .001 | 0.41 |
| Fusiform (L) | 11.8 | 3001.6 | 28.31 | < .001 | 0.39 |
| Pars Triangularis (L) | 12.5 | 3238.3 | 29.15 | < .001 | 0.38 |
| Caudal Anterior Cingulate (R) | 12.6 | 3340.5 | 29.04 | < .001 | 0.38 |
| Superior Temporal (R) | 11.9 | 3187.3 | 27.7 | < .001 | 0.37 |
| Inferior Parietal (L) | 6.8 | 1861.4 | 22.18 | < .001 | 0.36 |
| Rostral Anterior Cingulate (L) | 14.7 | 4033.6 | 34.91 | < .001 | 0.36 |
| Supramarginal (R) | 8.1 | 2334.5 | 21.94 | < .001 | 0.35 |
| Superior Frontal (L) | 8.4 | 2659.5 | 21.37 | < .001 | 0.32 |
| Parahippocampal (L) | 11.2 | 3629 | 25.65 | < .001 | 0.31 |
| Fusiform (R) | 9.5 | 3104.7 | 22.56 | < .001 | 0.31 |
| Insula (R) | 11.2 | 3837.3 | 26.04 | < .001 | 0.29 |
| Caudal Anterior Cingulate (L) | 9.8 | 3497.3 | 22.48 | < .001 | 0.28 |
| Supramarginal (L) | 5.6 | 2268.3 | 15.54 | < .001 | 0.25 |
| Transverse Temporal (R) | 9.3 | 3825.6 | 21.56 | < .001 | 0.24 |
| Lateral Occipital (L) | 6.3 | 2582.6 | 16.32 | < .001 | 0.24 |
| Precentral (L) | 8.6 | 3562.5 | 19.71 | < .001 | 0.24 |
| Lingual (R) | 4.9 | 2256.4 | 14.09 | < .001 | 0.21 |
| Lateral Occipital (R) | 5.0 | 2590.7 | 13.12 | < .001 | 0.19 |
| Pericalcarine (R) | 6.4 | 3379.6 | 14.76 | < .001 | 0.19 |
| Pars Orbitalis (R) | 5.9 | 3130.8 | 13.97 | < .001 | 0.19 |
| Banks of the Superior Temporal Sulcus (R) | 6.1 | 3335.7 | 14.1 | < .001 | 0.18 |
| Insula (L) | 7.0 | 4070.2 | 16.43 | < .001 | 0.17 |
| Paracentral (R) | 4.9 | 3022.5 | 11.68 | .001 | 0.16 |
| Rostral Anterior Cingulate (R) | 6.0 | 3723.7 | 13.68 | < .001 | 0.16 |
| Frontal Pole (R) | 5.9 | 3725.2 | 13.59 | < .001 | 0.16 |
| Postcentral (L) | 3.7 | 2434.1 | 10 | .002 | 0.15 |
| Superior Parietal (L) | 2.7 | 1827.6 | 8.9 | .003 | 0.15 |
| Posterior Cingulate (L) | 4.7 | 3215.4 | 10.97 | .001 | 0.15 |
| Inferior Parietal (R) | 2.6 | 1983.6 | 7.91 | .005 | 0.13 |
| Posterior Cingulate (R) | 3.8 | 3252.2 | 8.81 | .003 | 0.12 |
| Precuneus (L) | 1.8 | 1670.1 | 6.53 | .011 | 0.11 |
| Cuneus (R) | 2.3 | 2473.3 | 6.35 | .012 | 0.09 |
| Lateral Orbital Frontal (L) | 2.8 | 3221.5 | 6.49 | .011 | 0.09 |
| Parahippocampal (R) | 3.1 | 3619.3 | 7.08 | .008 | 0.09 |
| Medial Orbital Frontal (L) | 3.3 | 3909.8 | 7.58 | .006 | 0.08 |
| Superior Parietal (R) | 1.3 | 1910.9 | 4.19 | .041 | 0.07 |
| Postcentral (R) | 1.6 | 2273.3 | 4.39 | .036 | 0.07 |
| Lateral Orbital Frontal (R) | 2.6 | 3779.3 | 5.92 | .015 | 0.07 |
| Pars Orbitalis (L) | 2.1 | 3227.9 | 4.91 | .027 | 0.07 |
| Lingual (L) | 1.3 | 2392.1 | 3.7 | .054 | 0.06 |
| Transverse Temporal (L) | 1.6 | 3751.7 | 3.57 | .059 | 0.04 |
| Isthmus Cingulate (R) | 1.3 | 3390 | 2.98 | .084 | 0.04 |
| Precuneus (R) | 0.6 | 1709.8 | 2.29 | .130 | 0.04 |
| Pericalcarine (L) | 1.3 | 3386.5 | 2.94 | .086 | 0.04 |
| Medial Orbital Frontal (R) | 1.1 | 3312.6 | 2.53 | .112 | 0.03 |
| Temporal Pole (R) | 1.1 | 4499.1 | 3.78 | .052 | 0.02 |
| Isthmus Cingulate (L) | 0.2 | 3349.9 | 0.42 | .516 | 0.01 |
| Cuneus (L) | 0.1 | 2460.4 | 0.27 | .604 | 0.004 |
| Temporal Pole (L) | 0.2 | 4521.4 | 0.6 | .438 | 0.004 |
| Entorhinal (L) | 0.1 | 4455.5 | 0.34 | .557 | 0.002 |
| Paracentral (L) | 0.1 | 2976.4 | 0.17 | .683 | 0.002 |
| Entorhinal (R) | 0.1 | 4488.1 | 0.23 | .628 | 0.001 |

*Note*. Cortical regions are sorted vertically by the percentage of variance (% Variance) accounted for by the main effect of sex-at-birth within each analysis. Associations that passed false-discovery-rate (FDR) correction are shaded. *SS_B_ =* sum of squares (between). *SS_W_* = sum of squares (within). *F* = *F*-statistic. *p* = *p*-value.
